# RPDynaFlow: Generating RNA–protein Conformation Ensembles by Atomic Conditional Flow Matching

**DOI:** 10.64898/2026.08.28.747734

**Authors:** Yuntao Li, Kexin Lu

## Abstract

Conformation ensembles of biomolecules provide the basis for understanding structural transformations and drug design. Deep-learning generative models have advanced protein and small molecule ensemble generation, while RNA–protein complexes remain unaddressed due to the chemical heterogeneity, limited dataset size and the different flexibility scales of RNA and protein components. We present RPDynaFlow, a flow-matching model to generate conformation ensembles of RNA–protein complexes, trained on 600 ns trajectories of molecular dynamics (MD) simulation. The results show our model extends the sampling range of the phase space compared to MD simulation, which could be treated as a rapid and efficient complement to MD trajectories for studying RNA–protein interactions.

## 1 Introduction

RNA–protein complexes influence every stage of gene expression, including transcriptional regulation, pre-mRNA splicing, ribosomal translation and post-transcriptional silencing [1]. Instead of single structures, it is conformation ensembles that interpret the overall process of functioning. Riboswitches need large-scale rearrangements for ligand binding, spliceosomal snRNPs need coordinated domain motions to remodel RNA duplexes, and RNA-induced silencing complexes need to flex for accommodating diverse guide-target pairings [1]. Atomic resolution for these ensembles is essential for rational design of RNA-targeting therapeutics and studying RNA-binding proteins.

Molecular dynamics (MD) simulation is the physical standard when sampling biomolecular conformation, but still slow for exploring dynamic motions of RNA–protein complexes. Deeplearning generative models have recently demonstrated that meaningful conformation ensembles can be generated rapidly without long-time simulation. DiG [2] predicts equilibrium distributions by diffusion generating. BioEmu [3] was trained on *∼*200 ms of MD dataset to predict equilibrium distributions. AlphaFlow [4] applies flow matching to generate protein ensembles from sequence. SimpleFold [5] uses Boltzmann generators for single-chain proteins. PLACER [6] extends ensemble generation to protein–ligand complexes. These methods focus on either single-chain proteins, protein–protein systems, or protein–small-molecule pairs.

Existing generative methods have not focused on RNA–protein complexes, largely because of limited structural data, the demands to jointly model amino acid and nucleotide chemistry, and the wider flexibility range of RNA components, which requires finer scale.

We introduce RPDynaFlow(RNA–protein Dynamic Conformation Ensembles Flow-matching Generator), a flow-matching model trained on only 600ns of molecular dynamics trajectories. It generates all-atom conformation ensembles of RNA–protein complexes and extends the sampled phase space beyond the input simulations, serving as a fast and efficient complement to MD for studying RNA–protein interactions. In this article, we are also trying to explore the feasibility and performance of models trained under even more limited datasets.

## 2 Results

### 2.1 Model overview

RPDynaFlow learns per-atom displacement distributions around a static experimental structure using conditional flow matching. Given a single PDB structure, the model constructs an atomic graph connecting all heavy-atom pairs within 8 Å, then learns a velocity field that transports samples from a Gaussian prior to the target displacement distribution, as illustrated in Figure 1. At inference, Euler integration of the learned velocity field generates independent conformations, which can be run on a single GPU like RTX 4060Ti 8G.

**Figure 1:**
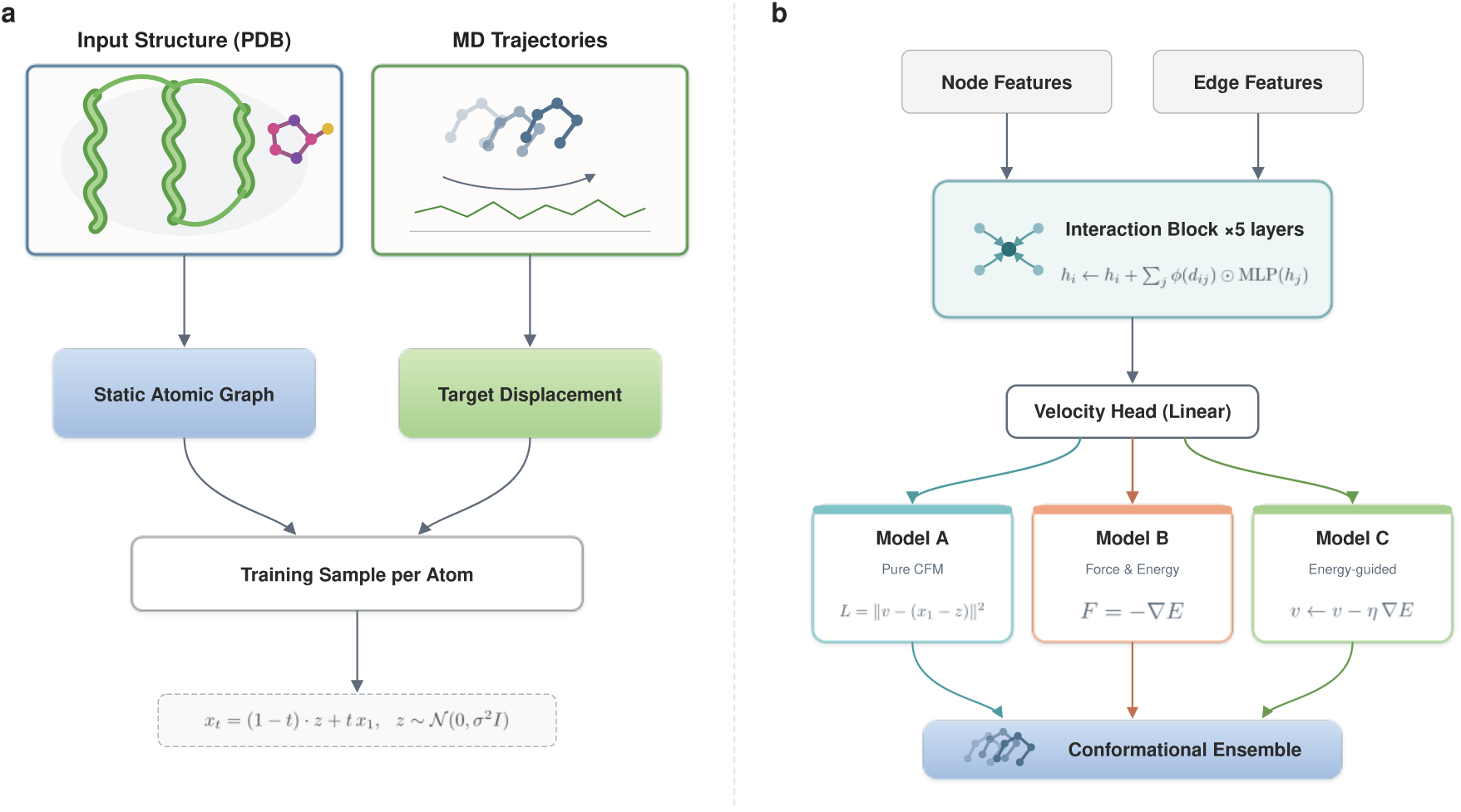
RPDynaFlow architecture and training scheme. **a.** The atomic graph was founded on input static structure and trajectories of 40ns MD per structure for conditional flow matching. **b.** The graph neural network processes atom velocities from node and edge features. We present 3 variants: Model A uses only the flow matching loss, Model B adds force and energy supervision, and Model C applies energy gradient guidance during sampling.

**Figure 2:**
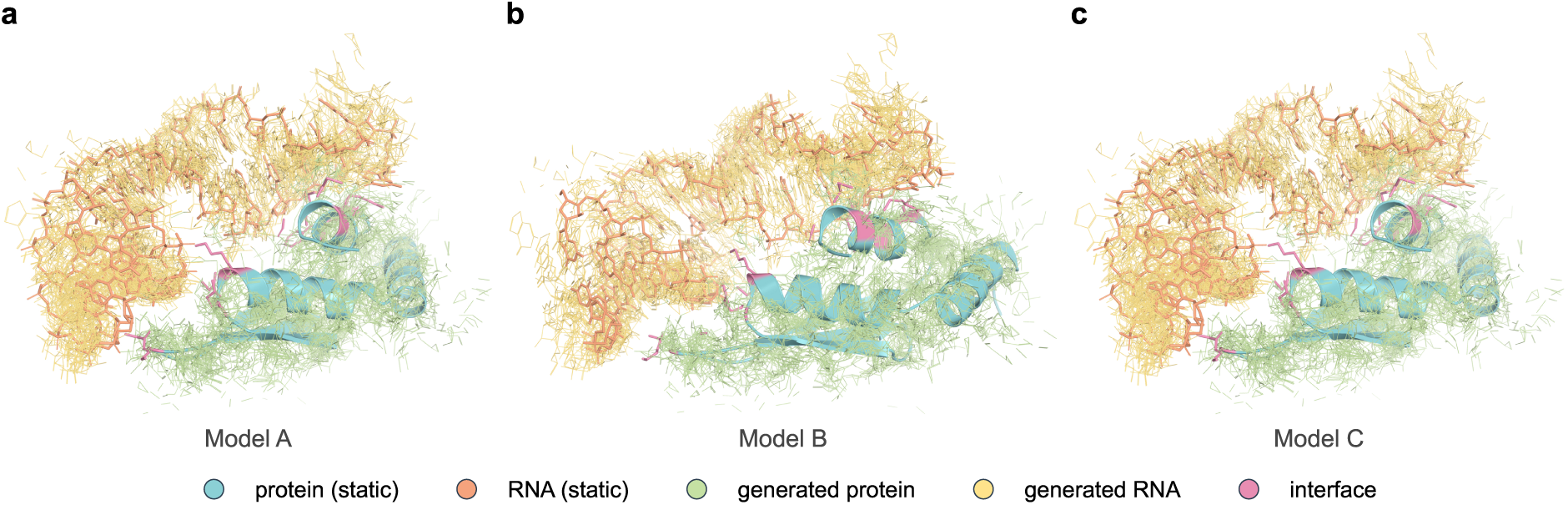
Structural visualization of generated ensembles for test system 2LBS. **a.** Model A generated ensemble (10 of 4,000 conformations). **b.** Model B generated ensemble. **c.** Model C generated ensemble. Protein is shown in teal/mint and RNA in orange/yellow; static structure underlaid for reference. All three models sample conformation diversity around the experimental structure, with B/C showing slightly larger displacement amplitudes.

The model is trained on short MD trajectories of 40 ns each from 15 small RNA–protein complexes, which contain between 1,144 and 3,444 heavy atoms as summarized in Table 2. Evaluation is performed on five held-out test systems including 1,368 to 2,258 heavy atoms that share no sequence homology with the training set, as shown in Supplementary Figure S1. For each test system, we generate 4,000 conformations and compare them against 401 MD reference frames sampled every 100 ps from their corresponding 40 ns trajectory.

**Table 1:** Summary of primary metrics averaged across five test systems. Arrows indicate preferred direction. Best value per metric in bold.

| Metric | Direction | Model A | Model B | Model C |
| --- | --- | --- | --- | --- |
| RMSF $r$ (complex) | ↑ | 0.586 | 0.646 | **0.649** |
| RMSF $r$ (protein) | ↑ | 0.664 | 0.738 | **0.742** |
| RMSF $r$ (RNA) | ↑ | 0.470 | **0.631** | 0.631 |
| RMWD (Å) | ↓ | **5.65** | 6.94 | 6.84 |
| PCA $W_2$ (Å) | ↓ | **5.47** | 6.78 | 6.70 |
| PC1 cosine sim | ↑ | 0.121 | 0.158 | **0.179** |
| IDDT (complex) | ↑ | **0.596** | 0.509 | 0.512 |
| ICS (generated) | ↑ | **0.730** | 0.639 | 0.638 |
| Weak Jaccard | ↑ | 0.440 | 0.505 | **0.510** |
| Transient Jaccard | ↑ | 0.139 | 0.164 | **0.175** |
| Bond MAE (Å) | ↓ | **0.276** | 0.423 | 0.380 |
| Chirality inversion rate (%) | ↓ | **2.6** | 11.0 | 7.5 |

**Table 2:** RNA–protein complexes used in this study. Heavy-atom counts exclude hydrogen, water, and buffer ions.

| PDB ID | Heavy atoms | Description | Role |
| --- | --- | --- | --- |
| 1NYB | 1,144 | RNA-binding domain-hairpin | train |
| 2ESE | 2,100 | RRM-stem-loop | train |
| 1EKZ | 2,173 | dsRBD-dsRNA | train |
| 1A1T | 1,514 | U1A-UTR hairpin | train |
| 4PDB | 1,831 | PUF-target RNA | train |
| 1DK1 | 1,978 | tRNA synthetase-tRNA | train |
| 2XDB | 2,055 | KH domain-polyC | train |
| 2Y8W | 2,098 | DEAD-box-ssRNA | train |
| 6GBM | 2,290 | Cas13-crRNA | train |
| 2FY1 | 2,343 | RNase III-dsRNA | train |
| 2N82 | 2,385 | Pumilio-PRE | train |
| 2L2K | 2,438 | Fox-1 RRM-UGCAUGU | train |
| 2N3O | 2,621 | Lin28-pre-let-7 | train |
| 1RKJ | 3,403 | SRP54-SRP RNA | train |
| 1FJE | 3,444 | Ribosomal L11-rRNA | train |
| 2LBS | 1,368 | FinO-stem-loop | test |
| 6TPH | 1,420 | SARS-CoV-2 Nsp9-RNA | test |
| 2HGH | 1,885 | Ro autoantigen-Y RNA | test |
| 7K9D | 1,990 | SARS-CoV-2 N-RNA | test |
| 4M4O | 2,258 | FUS RRM/ZnF-stem-loop | test |

To examine the effect of physics-based supervision, we introduce three model variants. Model A is trained with only the flow matching loss *L* = ∥*v_θ_ −* (*x*_1_ *− z*)∥^2^. Model B adds an auxiliary energy head with force and energy losses. Model C uses the weights of Model B but applies per-step energy gradient guidance. This allows us to isolate the contributions of explicit physical supervision from the generative objective.

### 2.2 Conformation space coverage

To assess how RPDynaFlow samples conformation space compared to MD, we performed joint principal component analysis (PCA) on the combined generated and reference ensembles, projecting onto the two leading principal components of the pooled C*α*/C3*^′^* coordinates (Figure 3).

**Figure 3:**
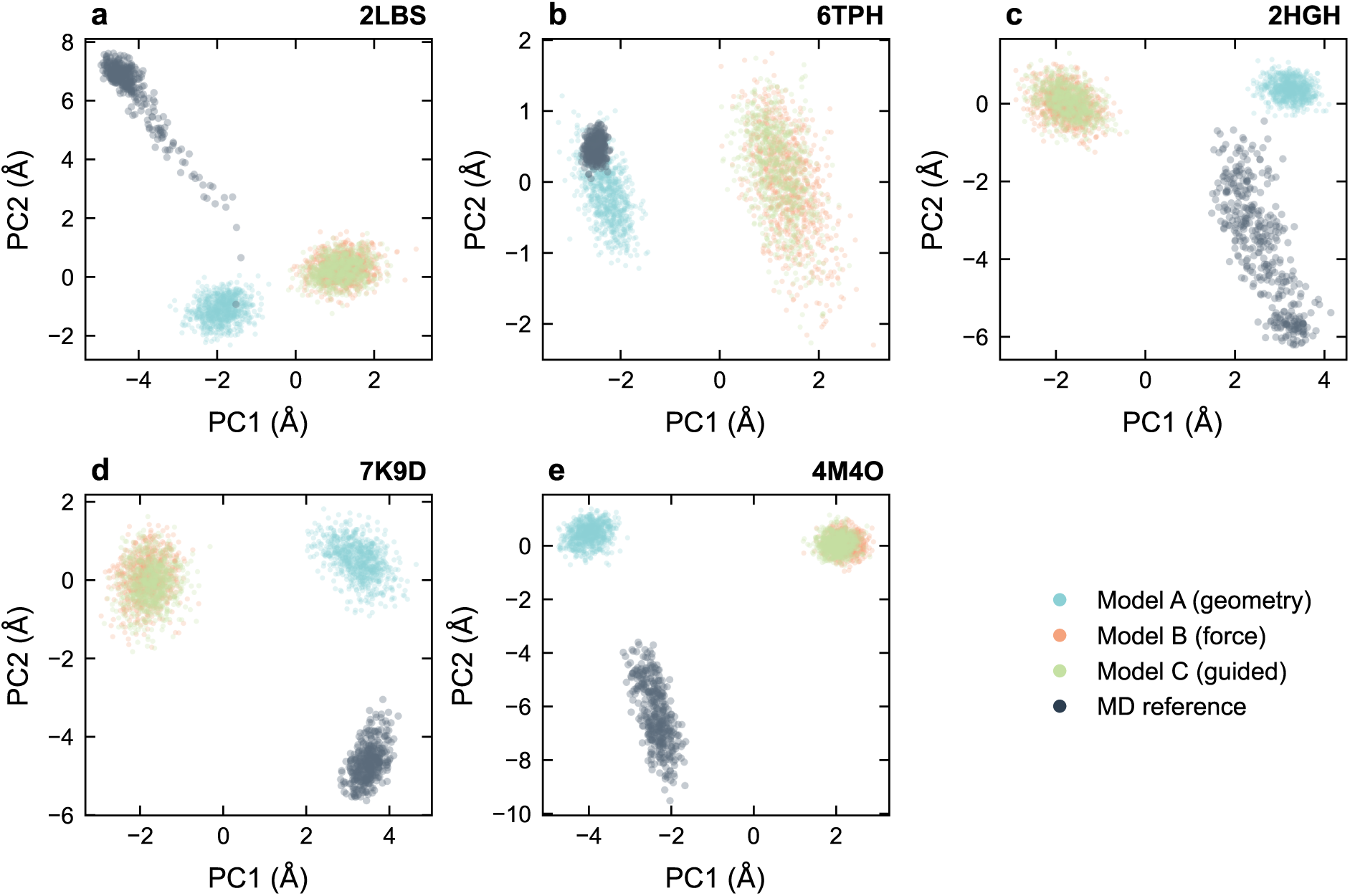
Generated ensembles cluster in the basin while MD shows large-amplitude motions . The PCA maps of the 5 test system. Different color spots shows individual conformations projected onto the first two principal components of the pooled C*α*/C3*^′^* coordinate set.

**Figure 4:**
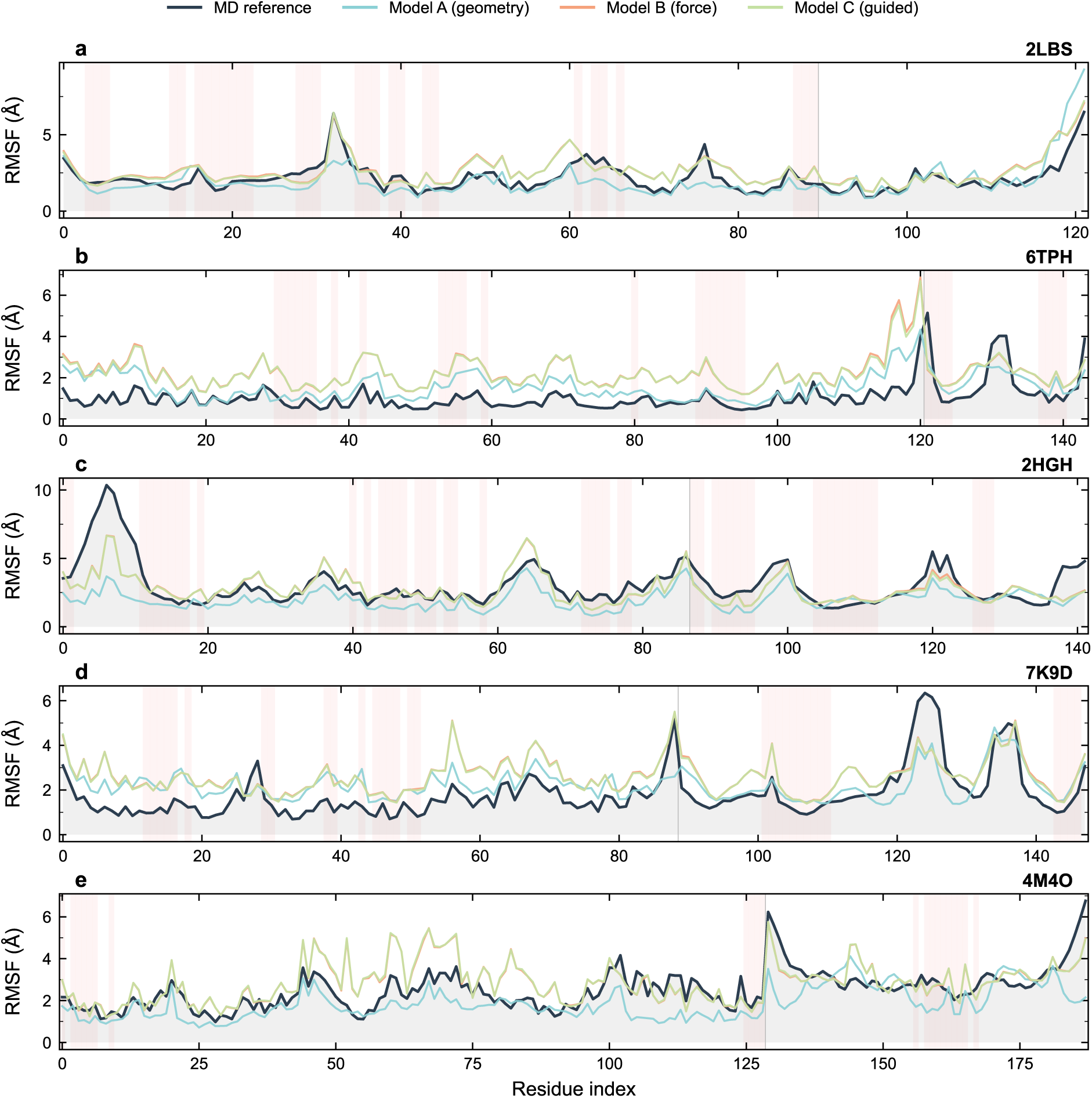
Force-supervised models improve flexibility reproduction. RMSF profiles for representative test systems are shown together with RMSF correlations across all systems, partitioned by complex, protein, RNA, and interface atoms. Models B and C are almost in a superposition, outperforming Model A, with the largest gains for RNA atoms.

**Figure 5:**
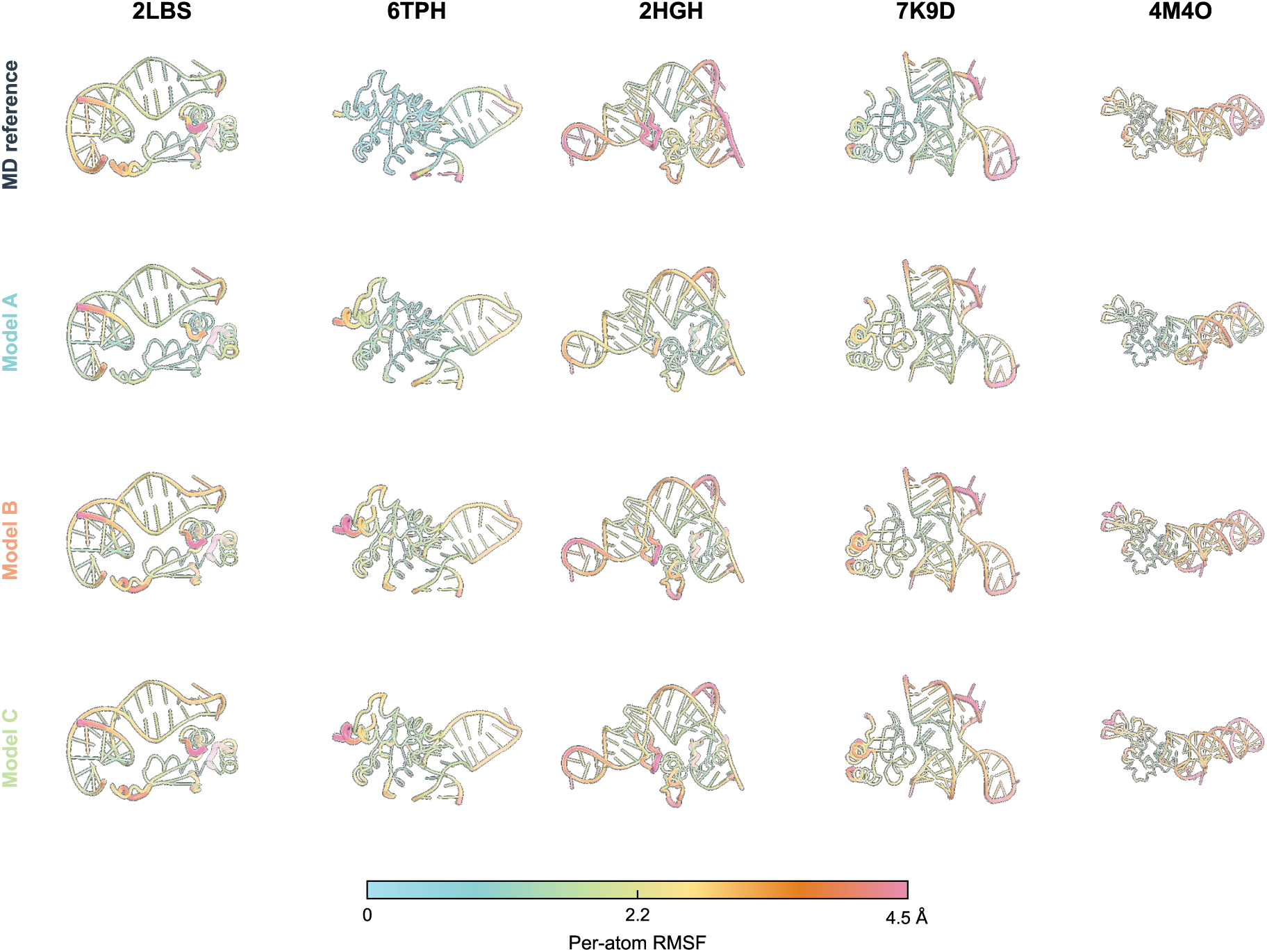
Mean-shift dominates distributional discrepancy. RMSF-coloured representations for all test systems, arranged by row for MD reference and Models A, B, and C. All generated models reproduce the spatial pattern of flexibility differing in amplitude.

Across all five test systems, the MD reference trajectories show progressive drift along PC2, reflecting large-amplitude repositioning. In contrast, the generated ensembles cluster near the starting structure with tighter but non-trivial spread. The Wasserstein-2 distance between the 2D PCA score distributions, approximated as Gaussians, averages 5.5 Å for Model A and 6.8 Å for Models B/C (Table 1), reflecting the larger displacement amplitudes learned by the force-supervised variants.

### 2.3 Flexibility patterns

Models B and C achieve consistently higher root-mean-square fluctuation (RMSF) correlations than Model A, with the largest improvement observed for RNA atoms, where the relative flexibility of loops and helices is better captured. Interface atoms remain the most challenging, showing the weakest RMSF correlations across all models. Despite the small training set and the added complexity of heteromolecular complexes, these correlations are broadly comparable to those reported by AlphaFlow[4] on single-chain proteins.

Distributional analysis further shows that the main discrepancy between generated and MD ensembles arises from mean displacement magnitude rather than variance structure. The root-mean Wasserstein distance is dominated by the mean-shift component, indicating that the models capture relative fluctuation amplitudes well but with shifted mean positions. Detailed per-system bond geometry is provided in Supplementary Figure S13.

### 2.4 Interface contact dynamics

The protein–RNA interface is the functionally critical region where binding specificity and allosteric communication are encoded. We characterize interface dynamics through per-contact distance distributions, interface contact similarity, and dynamic contact Jaccard indices.

Figure 6 shows per-contact-site heavy-atom distance distributions for the smallest test system 2LBS, where all interface contacts can be visualized. A clear pattern emerges: generated ensembles maintain tight, unimodal distance distributions centered near the crystallographic contact distance, whereas the MD reference shows bimodal or broadly distributed distances reflecting larger-amplitude structural changes over the 40 ns trajectory. This indicates that RPDynaFlow learns the range of fluctuations compatible with the starting interface geometry.

**Figure 6:**
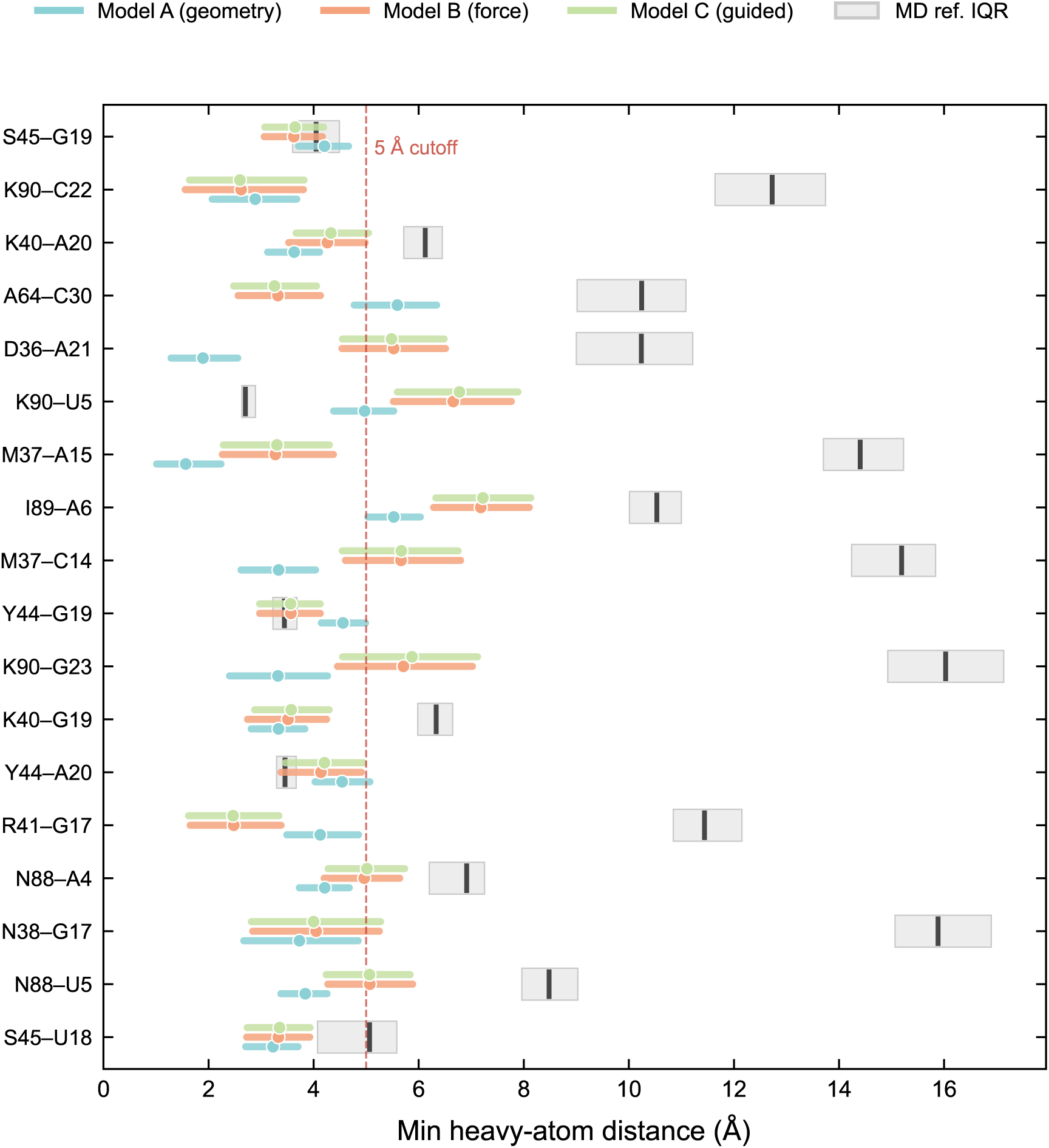
Interface contacts for 2LBS. The distribution of minimum heavy-atom distances for each RNA–protein residue pair across the ensemble. Filled distributions correspond to generated ensembles, and outlined distributions to the MD reference. The dashed line marks the 5 Å contact threshold.

Interface contact similarity (ICS) quantifies the overlap between static contacts and ensemble-averaged contacts. Generated ensembles achieve ICS values of 0.64–0.73, and the MD reference drops to 0.17–0.86 depending on the degree of structural rearrangement in each system. Contact occupancy maps are provided in Supplementary Figures S3–S7. The weak-contact Jaccard index for contacts present in more than 90% of frames averages 0.44 for Model A and 0.51 for Models B and C, indicating that the force-supervised variants better reproduce persistently maintained contacts. Transient contacts, defined as those present in more than 10% of frames, show Jaccard indices of 0.14–0.18, which reflects the difficulty of reproducing rare conformational states from limited training data.

### 2.5 RNA pose and internal deformation

We separate rigid-body RNA repositioning from internal RNA deformation using protein-anchored and RNA-anchored RMSD. Protein-anchored RMSD measures RNA displacement relative to the protein, while RNA-anchored RMSD captures internal conformation changes after superimposing the RNA onto itself, as shown in Figure 8.

**Figure 7:**
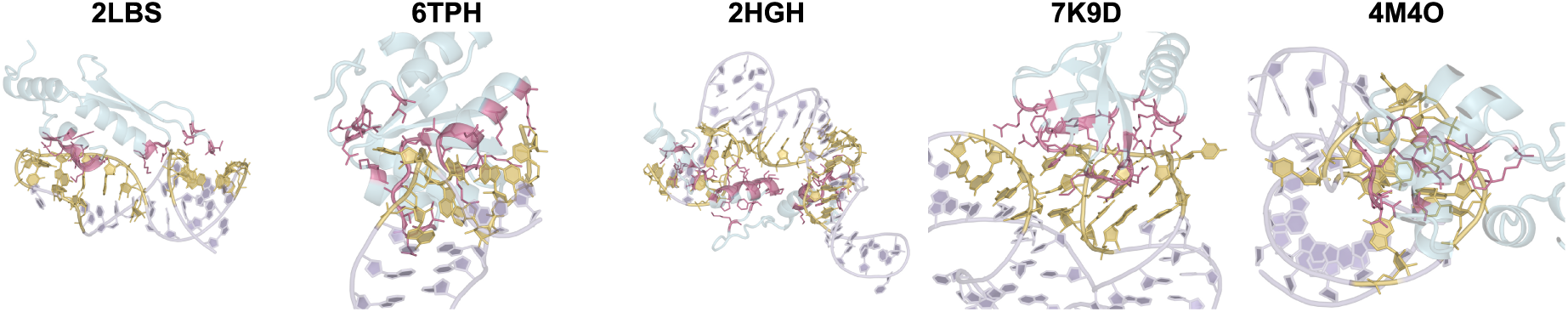
Interface metrics across test systems. Left: interface contact similarity for generated and MD reference ensembles. Right: weak and transient contact Jaccard indices. Higher values indicate better agreement with the static contact pattern. Models B and C consistently outperform Model A.

**Figure 8:**
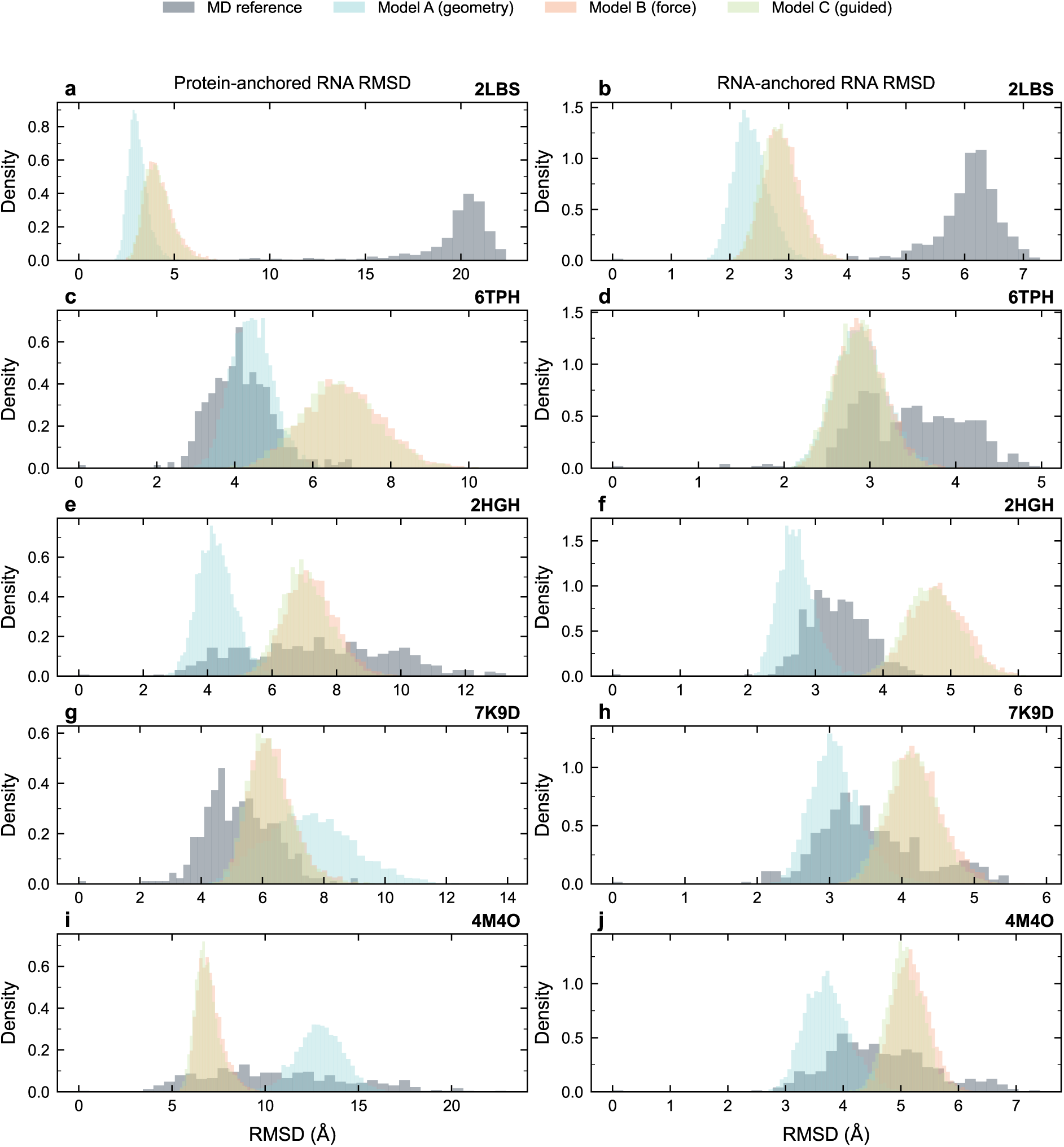
RNA pose and internal deformation. Left: protein-anchored RNA RMSD distributions. Right: RNA-anchored RNA RMSD distributions.

**Figure 9:**
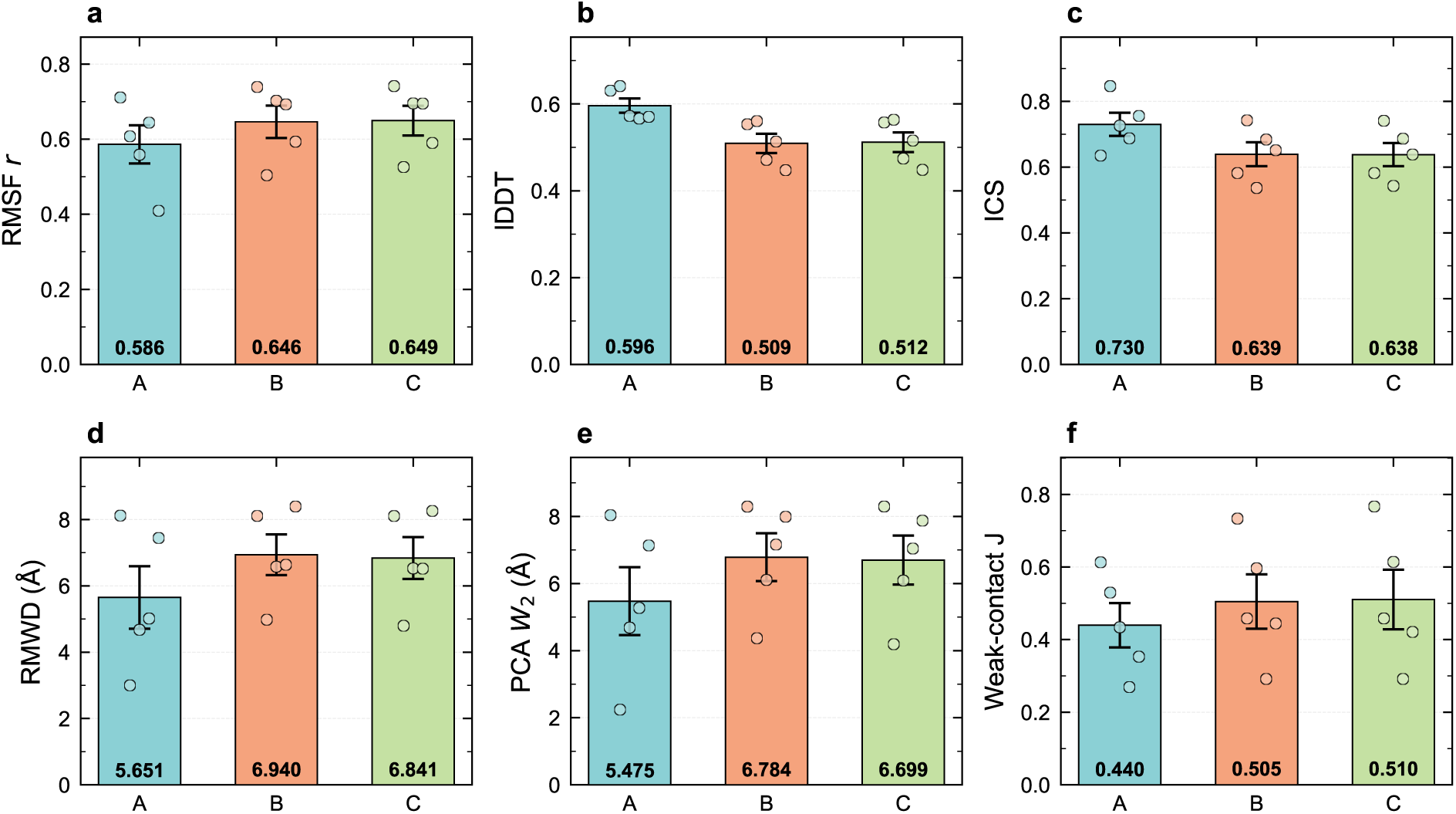
Model A leads structural fidelity; Models B and C lead flexibility. Per-system performance for each model variant across structural and flexibility metrics.

**Figure 10:**
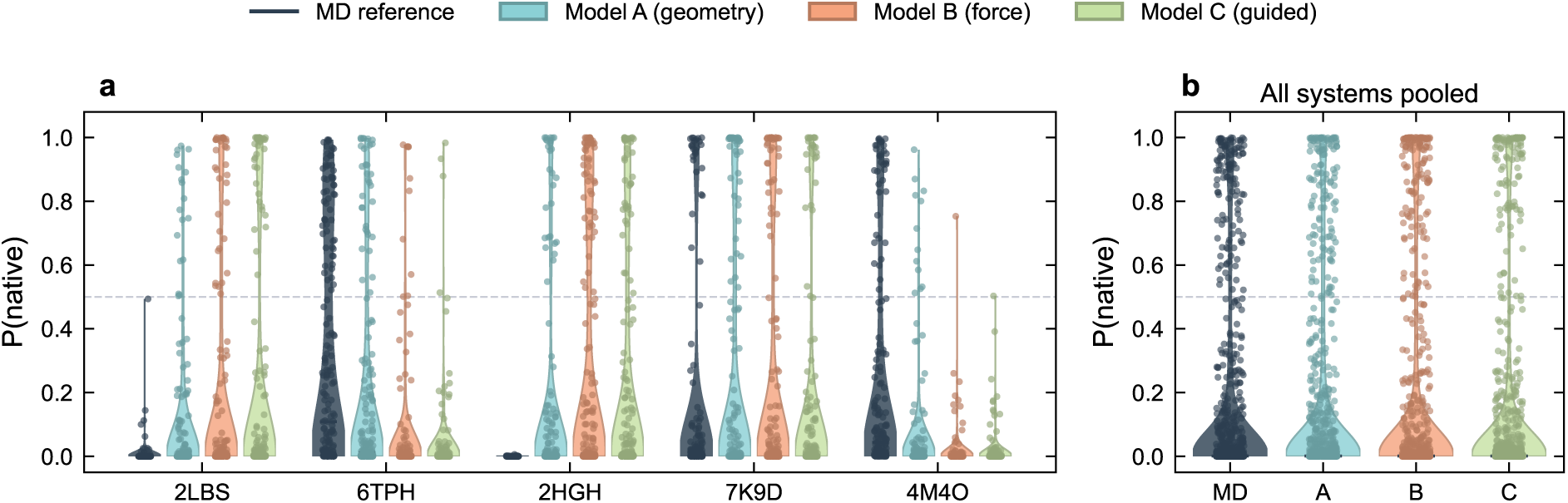
Generated and MD ensembles show comparable DRPScore distributions. DRPScore distributions for generated and MD frames are similar, indicating comparable structural plausibility.

The protein-anchored RMSD distributions of the MD reference are considerably broader than those of the generated ensembles, indicating that MD samples larger RNA displacements relative to the protein. In contrast, RNA-anchored RMSD distributions overlap substantially between generated and MD ensembles, demonstrating that the models reproduce internal RNA flexibility with similar amplitudes.

### 2.6 Structural fidelity and chemical validity

We evaluate structural fidelity using the local distance difference test (lDDT) [7]. Model A achieves the highest lDDT, followed by Models C and B, though all generated ensembles fall below the MD self-reference. The lDDT difference reflects diversity in the nature of the two distributions, as detailed in Supplementary Figure S12. Interchain lDDT shows the same ranking but lower values, confirming that the interface region remains more challenging to model than intra-chain contacts.

Chemical validity is assessed by bond-length mean absolute error and chirality inversion rates. Model A maintains near-ideal geometry, while Models B and C show slightly elevated bond-length errors, likely due to tension introduced by the force-matching objective. Chirality inversion rates remain low for all variants, with the higher rates in B and C concentrated in flexible loop regions rather than secondary-structure elements.

As an independent structural plausibility check, we score generated frames with DRPScore [8]. The generated ensembles achieve native-likeness scores comparable to the MD reference, confirming that RPDynaFlow preserves structurally plausible interfaces.

### 2.7 Model comparison and Pareto analysis

The three model variants define a clear Pareto frontier between structural fidelity and flexibility, as summarized in Table 1. Model A achieves the highest geometric accuracy (lDDT, bond MAE, RMWD), while Models B and C improve flexibility-related metrics substantially (RMSF correlation, contact Jaccard), at a modest cost in structural fidelity. Model C offers slightly better chemical validity than Model B without additional training burden.

This trade-off has a clear physical interpretation: force supervision encourages larger-amplitude and more physically motivated displacements that better match MD flexibility patterns, occasionally at the expense of equilibrium geometry. The choice among variants therefore depends on the intended application. Model A is preferable for tasks requiring high geometric precision, whereas Models B and C are better suited for studies where realistic dynamic behavior is the priority.

## 3 Discussion

We have demonstrated that conditional flow matching can generate physically meaningful conformation ensembles of RNA–protein complexes from a single static structure, trained on only 15 short MD trajectories totaling 600 ns. The generated ensembles capture conformation diversity relevant to RNA–protein interactions, as evidenced by tight interface distance distributions, preserved contact patterns, and realistic internal RNA deformation.

### Complementarity with MD

RPDynaFlow and MD sample different aspects of conformation space. The 40 ns MD trajectories used as training and reference data are finite-time trajectories that may include large-amplitude motions; they do not represent equilibrium ensembles. RPDynaFlow, by contrast, generates independent samples from the learned distribution without temporal ordering. Nevertheless, the MD reference captures fast local fluctuations that dominate conformation entropy, while slower large-scale rearrangements may be accessed through enhanced sampling protocols that can be combined with RPDynaFlow’s rapid generation.

### The A/B/C Pareto frontier

The systematic comparison of model variants reveals that physics-based supervision improves flexibility reproduction at the cost of structural precision. Force matching may encourage larger-amplitude motions that better reproduce MD flexibility patterns but occasionally distort local geometry. Energy guidance (Model C) partially increased chemical validity without additional training, suggesting the energy landscape learned by Model B contains useful information. Model A is preferred for applications requiring structural accuracy, such as docking pose evaluation and contact prediction, while Models B and C are better suited for flexibility characterization, including allosteric site identification.

### Limitations

First, the model generates independent samples with no temporal ordering; time-conditioned extensions could enable kinetic pathway prediction. Second, training on 15 short trajectories may limit generalization to systems with dynamics qualitatively different from the training set, and the fixed 8 Å graph may be unable to accommodate large rigid-body motions. Third, the model uses no sequence or evolutionary information explicitly, relying entirely on the static 3D structure; incorporating MSA features could improve predictions.

### Outlook

Time-conditioned flow matching could enable trajectory-like sampling with correct kinetic ordering while retaining the generative framework. Training on enhanced-sampling data, such as replica exchange or metadynamics, may access rare conformation states beyond the reach of short MD trajectories. The fast generation time makes RPDynaFlow suitable for high-throughput virtual screening pipelines. Finally, the atomic-graph architecture is general: extension to DNA–protein, protein–ligand, or multi-chain RNA complexes requires appropriate training data, and the fixed-graph assumption can be relaxed through periodic graph rebuilding for applications involving large conformation changes.

## 4 Methods

### 4.1 Molecular systems and data

We selected 20 RNA–protein complexes from the RPpocket database [9], as listed in Table 2. The dataset comprises 15 training systems and 5 test systems. Selection criteria require a resolution of 3.5 Å or better, the presence of both protein and RNA chains in the asymmetric unit, a complex size between 1,000 and 3,500 heavy atoms, and the absence of sequence homology between training and test sets, verified by pairwise BLAST and shown in Supplementary Figure S1.

### 4.2 Molecular dynamics simulations

All-atom MD simulations were performed with GROMACS 2025.2 [10] using the CHARMM36 force field [11] for proteins and CHARMM36 for nucleic acids, with the TIP3P water model [12]. Each system was solvated in a rhombic dodecahedral box with a minimum buffer distance of 12 Å, neutralized with Na^+^ and Cl*^−^* ions at 150 mM. Following steepest-descent energy minimization to a tolerance of 1000 kJ*/*mol*/*nm, systems were equilibrated for 100 ps in the NVT ensemble and 100 ps in the NPT ensemble with position restraints on heavy atoms. Production runs of 40 ns used a 2 fs time step, LINCS constraints on hydrogen bonds [13], PME electrostatics with a 1.2 nm cutoff [14], the velocity-rescale thermostat at 300 K [15], and the Parrinello-Rahman barostat at 1 bar [16].

Post-processing included PBC correction, centering, and least-squares fitting to the initial frame using gmx trjconv -pbc cluster -center -ur compact -fit rot+trans. Coordinates were saved every 100 ps, yielding 401 frames per system, and only solute heavy atoms were retained, excluding hydrogen, water, and ions.

### 4.3 Feature representation

Each system is represented by a set of *A* heavy atoms. Each atom carries a coordinate vector *r ∈* R*^A×^*^3^ in angstroms, an element class *e_i_* among six categories, a residue type *t_j_* covering the 20 standard amino acids and four RNA nucleotides, and a residue index *j*(*i*) mapping the atom to its parent residue. The static structure *s* corresponds to the first frame, i.e., the experimental coordinates after alignment. Target displacements are scaled as *x*_1_ = (*r − s*)*/S* with *S* = 10 Å.

### 4.4 Conditional flow matching

We adopt the optimal-transport conditional flow matching framework [17, 18]. The forward process interpolates between noise and target as

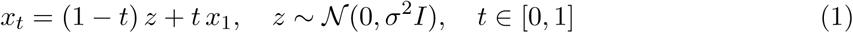

where *σ* = 0.368 is the per-dataset displacement standard deviation computed at training start. The model *v_θ_*(*x_t_, s, t*) is trained to predict the conditional velocity field defined by the loss

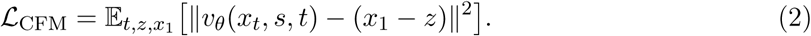

### 4.5 AtomFlowNet architecture

The neural network follows a SchNet-style message-passing architecture [19] operating on a fixed atomic graph. Edges connect atom pairs within 8 Å of the static structure and remain unchanged across all time steps and training frames. Inter-atomic distances are expanded into 16 Gaussian radial basis functions with learned centers and widths, multiplied by a cosine cutoff envelope. Each atom is described by the concatenation of its noised position *x*^(^*^i^*^)^, static position *s*^(^*^i^*^)^, element embedding with *d* = 16, residue-type embedding with *d* = 16, and sinusoidal time embedding with *d* = 16, projected to *d* = 128 dimensions. Five SchNet interaction blocks update node features through residual connections according to

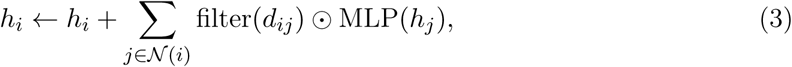

where filter maps edge features to filter weights. The model contains approximately 0.3 million parameters, and a final linear head maps node features to per-atom 3D velocity predictions.

### 4.6 Model B: force and energy supervision

Model B augments the conditional flow matching loss with physics-based auxiliary objectives. An energy head computes per-atom energies *e_i_* = MLP(*h_i_*), which are summed to a total energy 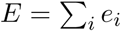 Forces are obtained by automatic differentiation as *F* = *−∇_x_E*. The combined loss is

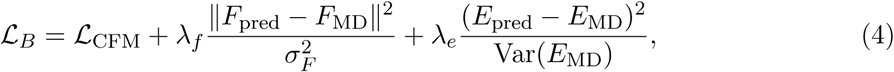

with *λ_f_* = 1.0 and *λ_e_* = 0.1. MD forces and energies are extracted from GROMACS energy files and mapped to the heavy-atom representation.

### 4.7 Model C: energy-guided sampling

Model C uses the trained weights of Model B without modification. During Euler integration, each velocity prediction is adjusted by the negative energy gradient as

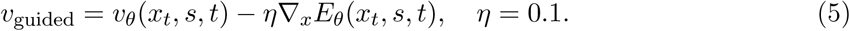

This steers sampling toward lower-energy regions of the learned potential surface without additional training.

### 4.8 Training details

All models were trained with AdamW [20], a learning rate of 3 *×* 10*^−^*^4^, weight decay of 10*^−^*^4^, and a cosine annealing schedule. Model A was trained for 150 epochs, while Model B required 300 epochs due to the auxiliary loss landscape. Each training step samples one system uniformly and eight random frames from that system’s trajectory. Training was performed on a single NVIDIA RTX 4060 Ti with 16 GB of memory; Model A completes in approximately 2.5 hours.

### 4.9 Evaluation framework

The evaluation framework adapts metrics from AlphaFlow [4], SimpleFold [5], and PLACER [6], all computed on GPU for efficiency. The local distance difference test (lDDT) is computed with an inclusion radius of 15 Å and thresholds of 0.5, 1.0, 2.0, and 4.0 Å, reported as the median over the generated ensemble and partitioned into complex, protein, RNA, and interchain subsets. Per-atom RMSF is calculated as the standard deviation of atomic positions across the ensemble, and Pearson correlation with MD RMSF profiles is evaluated by molecular component. The root-mean Wasserstein distance (RMWD) between per-atom one-dimensional displacement distributions is computed under Gaussian approximation and decomposed as 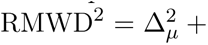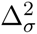.

Joint PCA is performed on pooled C*α* and C3*^′^* coordinates from the combined generated and MD ensembles, and the two-dimensional Gaussian Wasserstein-2 distance is computed between score distributions. Interface contacts are defined at a 5 Å heavy-atom minimum distance between residues following the OpenStructure convention [21]. Weak contacts are present in more than 90% of frames, while transient contacts appear in more than 10%. The Jaccard index between generated and MD contact sets is reported. Interface contact similarity is defined as ICS = 2*|C_S_ ∩ C_X_|/*(*|C_S_|* + *|C_X_|*), where *C_S_* denotes static contacts and *C_X_* per-frame contacts.

Bond-length mean absolute error is computed against ideal Engh-Huber distances [22], and chirality is assessed as the fraction of C*α* atoms with inverted stereocenters. Anchored RMSD separates pose change from internal deformation: protein-anchored RMSD superimposes protein C*α* atoms and measures RNA heavy-atom RMSD, while RNA-anchored RMSD superimposes RNA P/C3*^′^*/C1*^′^* atoms and measures RNA heavy-atom RMSD. DRPScore [8] provides an independent structural plausibility assessment using a 4D-CNN classifier trained on crystallographic interfaces. We scored 200 randomly selected frames per system and model, with preprocessing performed using BioPython-based voxelization. All per-system metrics are accompanied by 95% bootstrap confidence intervals based on 1,000 resamples. Evaluation code is available at https://github.com/YuntaoOvO/RPDynaFlow.

## Data and code availability

All code, trained checkpoints, evaluation scripts, and example data are available at https://github.com/YuntaoOvO/RPDynaFlow. MD trajectories and processed arrays are available upon request.

## Acknowledgements

This work was developed during the 2026 PEBBLE BioFusion Workshop at Westlake University, Hangzhou, China. We thank the workshop organizers, teaching assistants, and participants for stimulating discussions.

## Supplementary Information

**Table S1:**
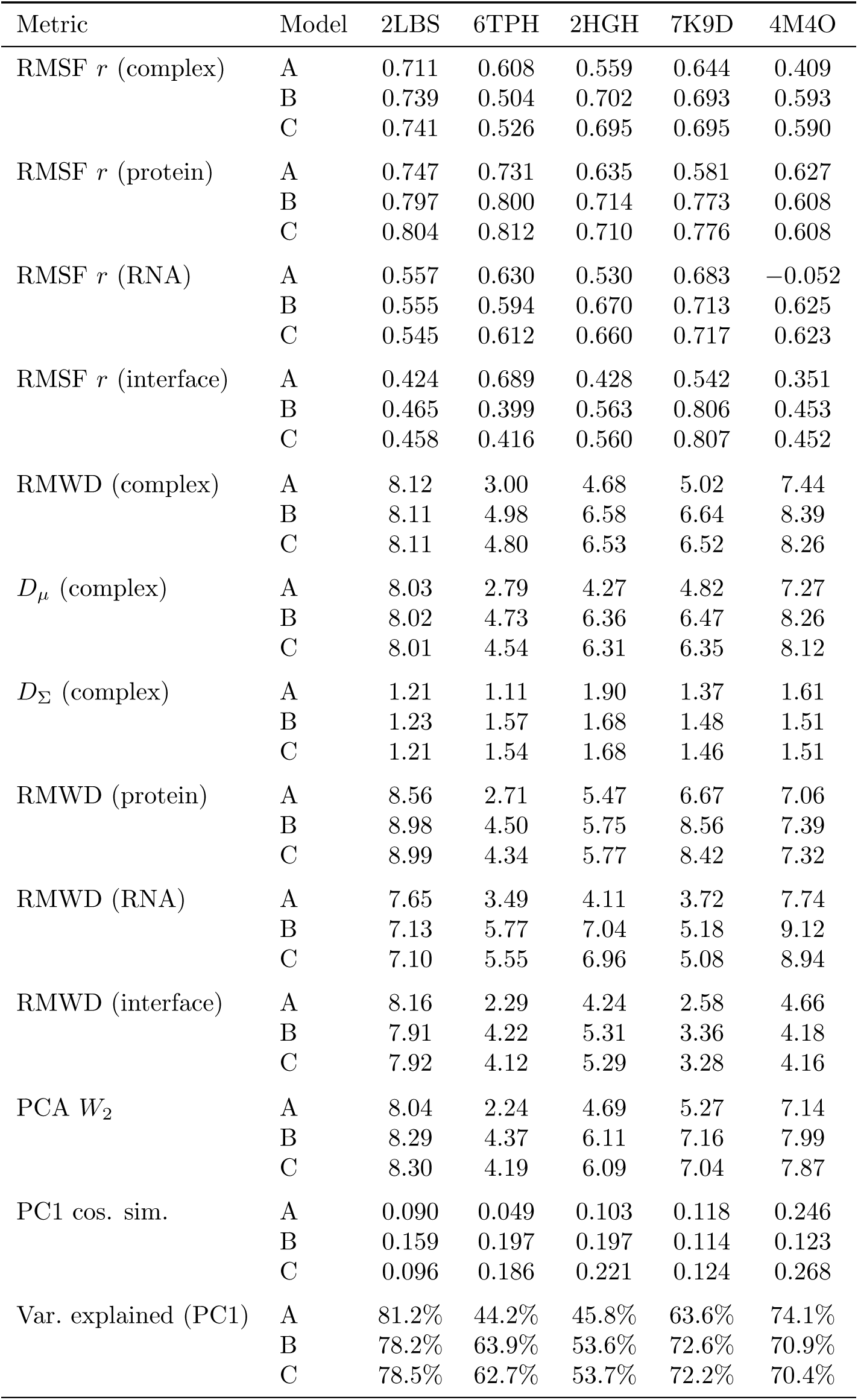
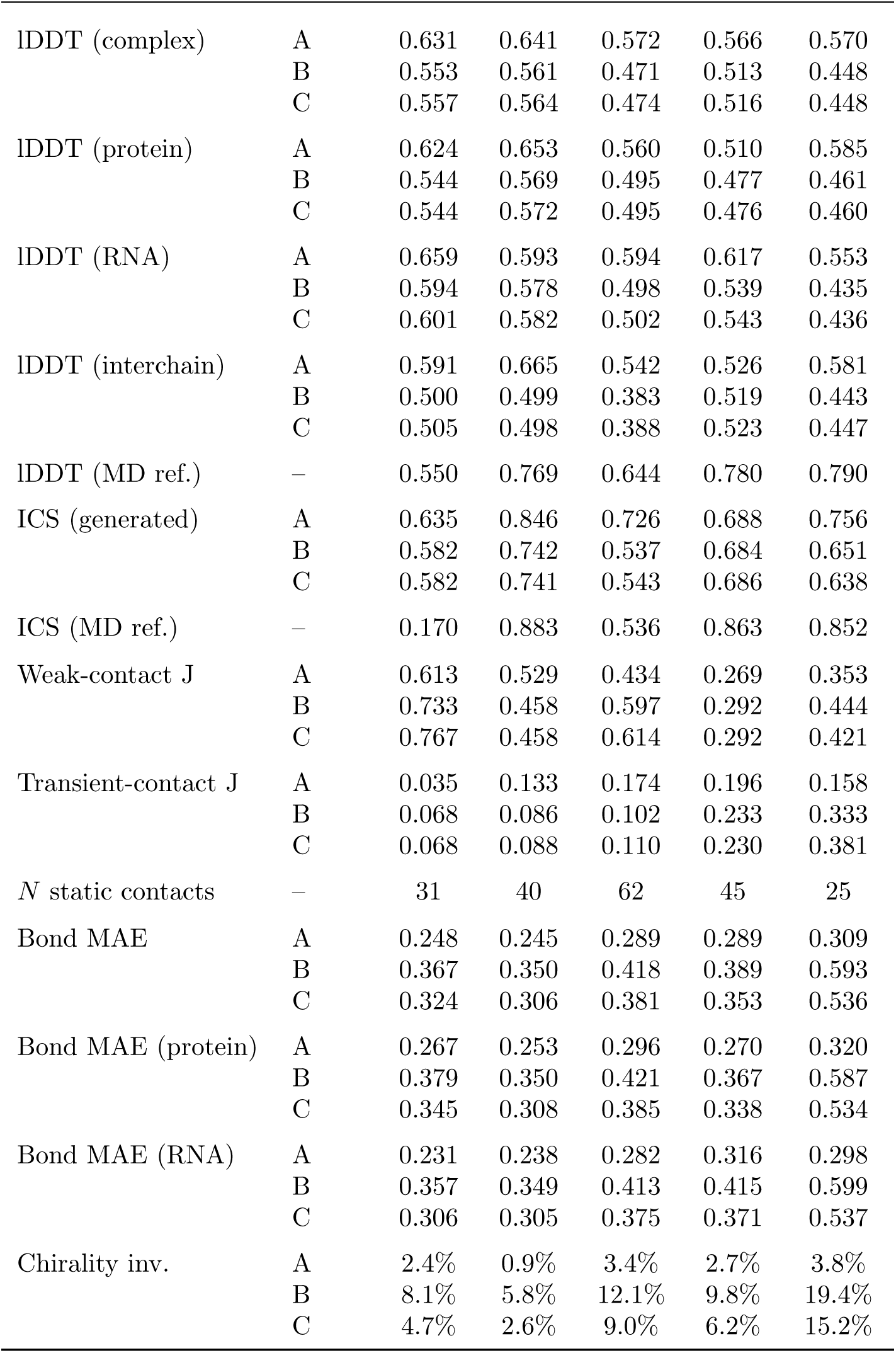
Complete primary evaluation metrics for all test systems and model variants. RMSF *r* = Pearson correlation (higher is better); RMWD = Root Mean Wasserstein Distance in Å (lower is better); PCA *W_2_* in Å (lower is better); lDDT = local Distance Difference Test (higher is better); ICS = Interface Contact Similarity (higher is better); Jaccard = set overlap (higher is better); Bond MAE in Å (lower is better).

| Metric | Model | 2LBS | 6TPH | 2HGH | 7K9D | 4M4O |
| --- | --- | --- | --- | --- | --- | --- |
| RMSF $r$ (complex) | A | 0.711 | 0.608 | 0.559 | 0.644 | 0.409 |
|  | B | 0.739 | 0.504 | 0.702 | 0.693 | 0.593 |
|  | C | 0.741 | 0.526 | 0.695 | 0.695 | 0.590 |
| RMSF $r$ (protein) | A | 0.747 | 0.731 | 0.635 | 0.581 | 0.627 |
|  | B | 0.797 | 0.800 | 0.714 | 0.773 | 0.608 |
|  | C | 0.804 | 0.812 | 0.710 | 0.776 | 0.608 |
| RMSF $r$ (RNA) | A | 0.557 | 0.630 | 0.530 | 0.683 | −0.052 |
|  | B | 0.555 | 0.594 | 0.670 | 0.713 | 0.625 |
|  | C | 0.545 | 0.612 | 0.660 | 0.717 | 0.623 |
| RMSF $r$ (interface) | A | 0.424 | 0.689 | 0.428 | 0.542 | 0.351 |
|  | B | 0.465 | 0.399 | 0.563 | 0.806 | 0.453 |
|  | C | 0.458 | 0.416 | 0.560 | 0.807 | 0.452 |
| RMWD (complex) | A | 8.12 | 3.00 | 4.68 | 5.02 | 7.44 |
|  | B | 8.11 | 4.98 | 6.58 | 6.64 | 8.39 |
|  | C | 8.11 | 4.80 | 6.53 | 6.52 | 8.26 |
| $D_\mu$ (complex) | A | 8.03 | 2.79 | 4.27 | 4.82 | 7.27 |
|  | B | 8.02 | 4.73 | 6.36 | 6.47 | 8.26 |
|  | C | 8.01 | 4.54 | 6.31 | 6.35 | 8.12 |
| $D_\Sigma$ (complex) | A | 1.21 | 1.11 | 1.90 | 1.37 | 1.61 |
|  | B | 1.23 | 1.57 | 1.68 | 1.48 | 1.51 |
|  | C | 1.21 | 1.54 | 1.68 | 1.46 | 1.51 |
| RMWD (protein) | A | 8.56 | 2.71 | 5.47 | 6.67 | 7.06 |
|  | B | 8.98 | 4.50 | 5.75 | 8.56 | 7.39 |
|  | C | 8.99 | 4.34 | 5.77 | 8.42 | 7.32 |
| RMWD (RNA) | A | 7.65 | 3.49 | 4.11 | 3.72 | 7.74 |
|  | B | 7.13 | 5.77 | 7.04 | 5.18 | 9.12 |
|  | C | 7.10 | 5.55 | 6.96 | 5.08 | 8.94 |
| RMWD (interface) | A | 8.16 | 2.29 | 4.24 | 2.58 | 4.66 |
|  | B | 7.91 | 4.22 | 5.31 | 3.36 | 4.18 |
|  | C | 7.92 | 4.12 | 5.29 | 3.28 | 4.16 |
| PCA $W_2$ | A | 8.04 | 2.24 | 4.69 | 5.27 | 7.14 |
|  | B | 8.29 | 4.37 | 6.11 | 7.16 | 7.99 |
|  | C | 8.30 | 4.19 | 6.09 | 7.04 | 7.87 |
| PC1 cos. sim. | A | 0.090 | 0.049 | 0.103 | 0.118 | 0.246 |
|  | B | 0.159 | 0.197 | 0.197 | 0.114 | 0.123 |
|  | C | 0.096 | 0.186 | 0.221 | 0.124 | 0.268 |
| Var. explained (PC1) | A | 81.2% | 44.2% | 45.8% | 63.6% | 74.1% |
|  | B | 78.2% | 63.9% | 53.6% | 72.6% | 70.9% |
|  | C | 78.5% | 62.7% | 53.7% | 72.2% | 70.4% |
| IDDT (complex) | A | 0.631 | 0.641 | 0.572 | 0.566 | 0.570 |
|  | B | 0.553 | 0.561 | 0.471 | 0.513 | 0.448 |
|  | C | 0.557 | 0.564 | 0.474 | 0.516 | 0.448 |
| IDDT (protein) | A | 0.624 | 0.653 | 0.560 | 0.510 | 0.585 |
|  | B | 0.544 | 0.569 | 0.495 | 0.477 | 0.461 |
|  | C | 0.544 | 0.572 | 0.495 | 0.476 | 0.460 |
| IDDT (RNA) | A | 0.659 | 0.593 | 0.594 | 0.617 | 0.553 |
|  | B | 0.594 | 0.578 | 0.498 | 0.539 | 0.435 |
|  | C | 0.601 | 0.582 | 0.502 | 0.543 | 0.436 |
| IDDT (interchain) | A | 0.591 | 0.665 | 0.542 | 0.526 | 0.581 |
|  | B | 0.500 | 0.499 | 0.383 | 0.519 | 0.443 |
|  | C | 0.505 | 0.498 | 0.388 | 0.523 | 0.447 |
| IDDT (MD ref.) | – | 0.550 | 0.769 | 0.644 | 0.780 | 0.790 |
| ICS (generated) | A | 0.635 | 0.846 | 0.726 | 0.688 | 0.756 |
|  | B | 0.582 | 0.742 | 0.537 | 0.684 | 0.651 |
|  | C | 0.582 | 0.741 | 0.543 | 0.686 | 0.638 |
| ICS (MD ref.) | – | 0.170 | 0.883 | 0.536 | 0.863 | 0.852 |
| Weak-contact J | A | 0.613 | 0.529 | 0.434 | 0.269 | 0.353 |
|  | B | 0.733 | 0.458 | 0.597 | 0.292 | 0.444 |
|  | C | 0.767 | 0.458 | 0.614 | 0.292 | 0.421 |
| Transient-contact J | A | 0.035 | 0.133 | 0.174 | 0.196 | 0.158 |
|  | B | 0.068 | 0.086 | 0.102 | 0.233 | 0.333 |
|  | C | 0.068 | 0.088 | 0.110 | 0.230 | 0.381 |
| *N* static contacts | – | 31 | 40 | 62 | 45 | 25 |
| Bond MAE | A | 0.248 | 0.245 | 0.289 | 0.289 | 0.309 |
|  | B | 0.367 | 0.350 | 0.418 | 0.389 | 0.593 |
|  | C | 0.324 | 0.306 | 0.381 | 0.353 | 0.536 |
| Bond MAE (protein) | A | 0.267 | 0.253 | 0.296 | 0.270 | 0.320 |
|  | B | 0.379 | 0.350 | 0.421 | 0.367 | 0.587 |
|  | C | 0.345 | 0.308 | 0.385 | 0.338 | 0.534 |
| Bond MAE (RNA) | A | 0.231 | 0.238 | 0.282 | 0.316 | 0.298 |
|  | B | 0.357 | 0.349 | 0.413 | 0.415 | 0.599 |
|  | C | 0.306 | 0.305 | 0.375 | 0.371 | 0.537 |
| Chirality inv. | A | 2.4% | 0.9% | 3.4% | 2.7% | 3.8% |
|  | B | 8.1% | 5.8% | 12.1% | 9.8% | 19.4% |
|  | C | 4.7% | 2.6% | 9.0% | 6.2% | 15.2% |

**Figure S1:**
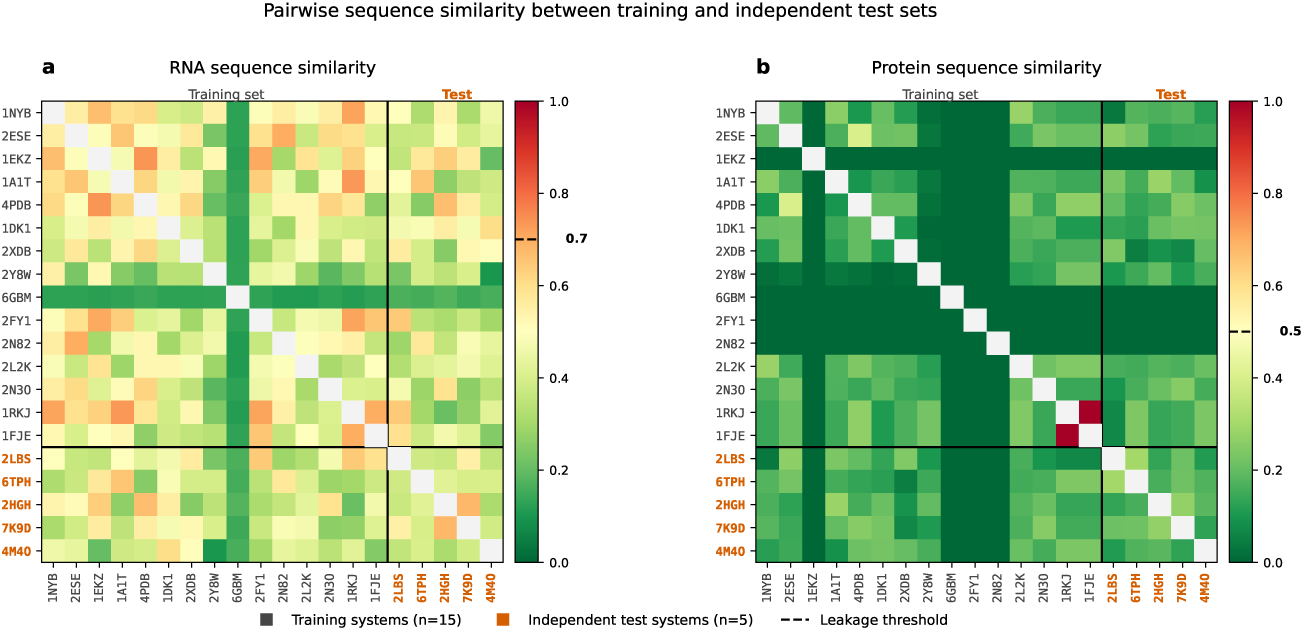
Sequence similarity between training and test systems. Pairwise protein sequence identity is shown in the upper triangle and RNA sequence identity in the lower triangle. No test system shares more than 25% sequence identity with any training system, supporting generalization rather than memorization.

**Figure S2:**
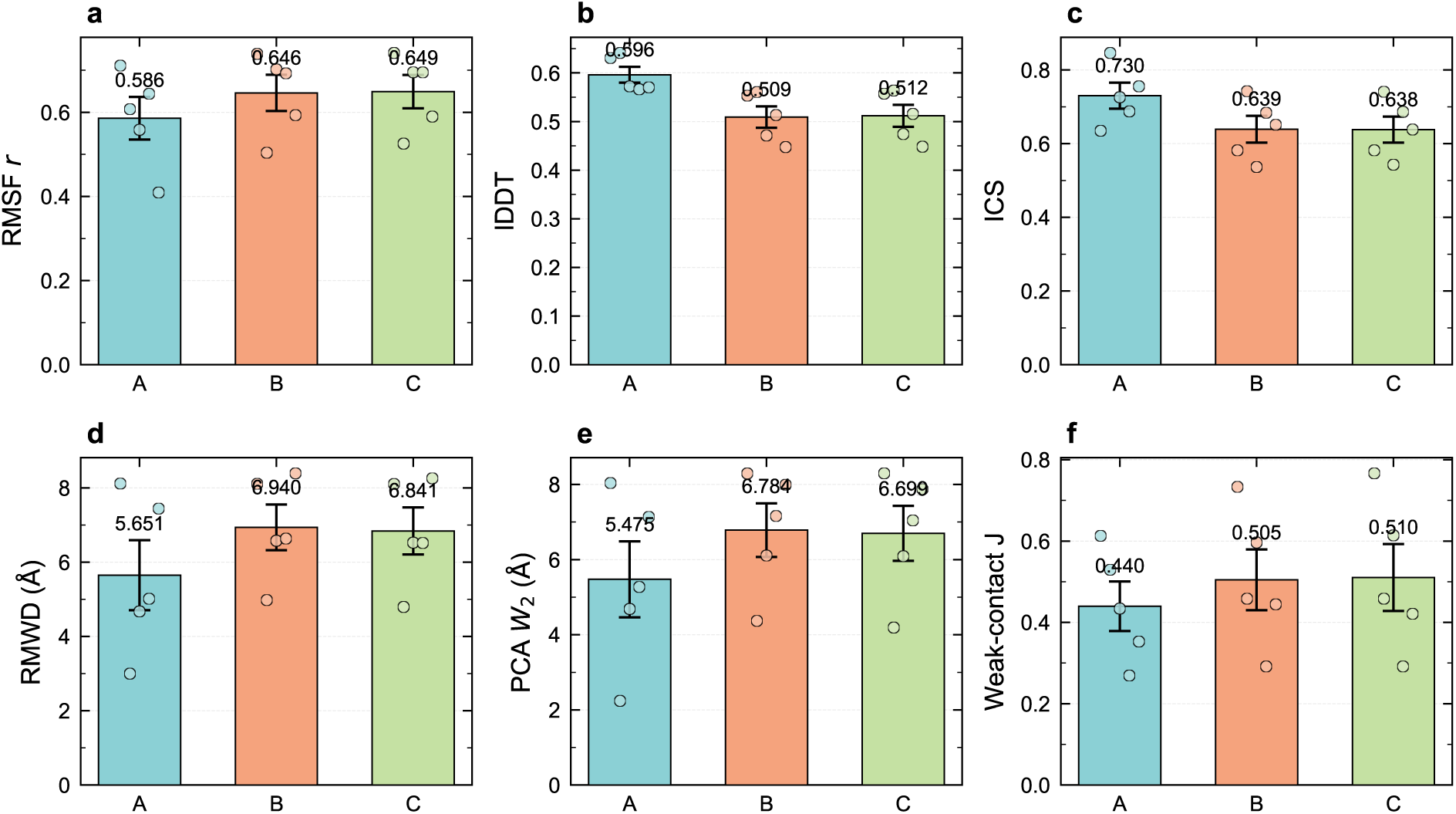
Per-system metric breakdown for all model variants. Values for the primary metrics are shown individually for each test system. Each bar group corresponds to Models A, B, and C.

**Figure S3:**
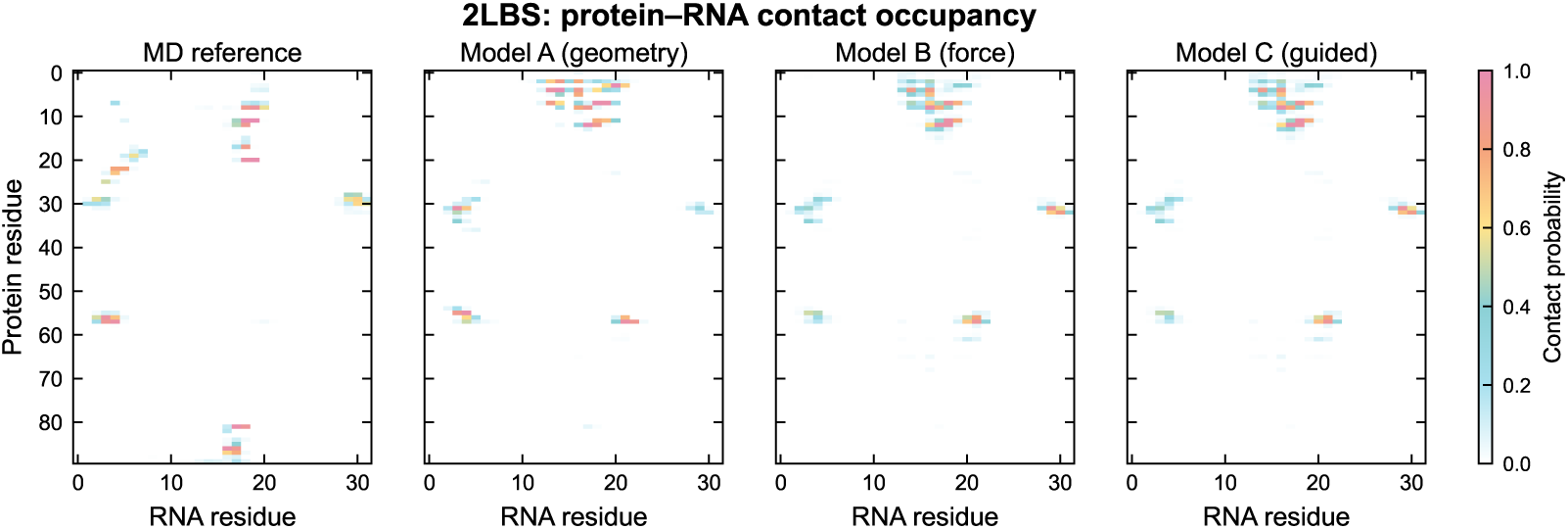
RNA–protein contact occupancy map for 2LBS. The heatmap shows the fraction of ensemble frames in which each RNA–protein residue pair is in contact, defined as a heavy-atom distance below 5 Å. Panels from left to right correspond to MD reference and Models A, B, and C.

**Figure S4:**
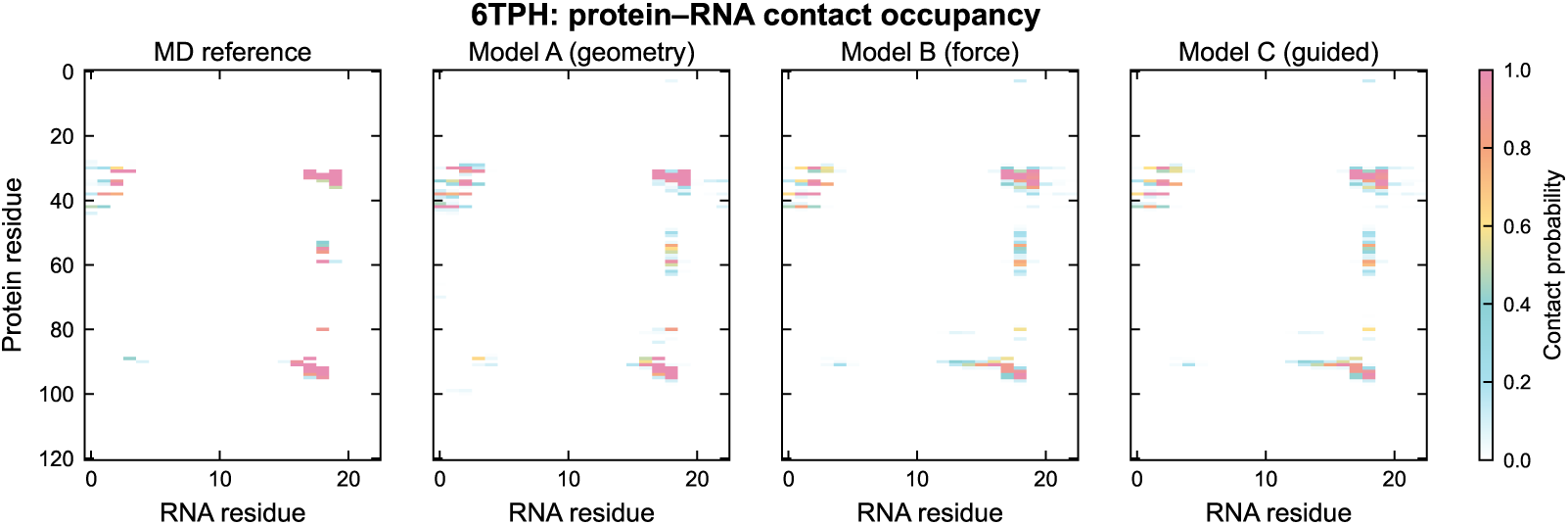
RNA–protein contact occupancy map for 6TPH. Layout follows the previous figure.

**Figure S5:**
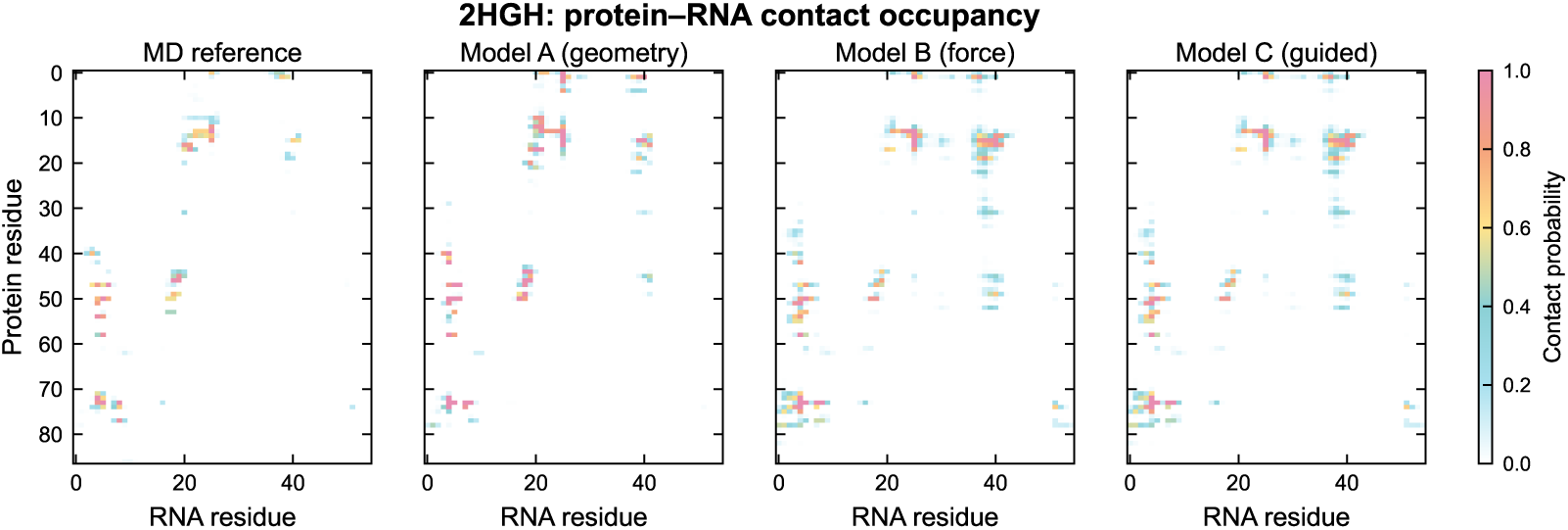
RNA–protein contact occupancy map for 2HGH. Contact patterns differ between the MD reference and the generated ensembles.

**Figure S6:**
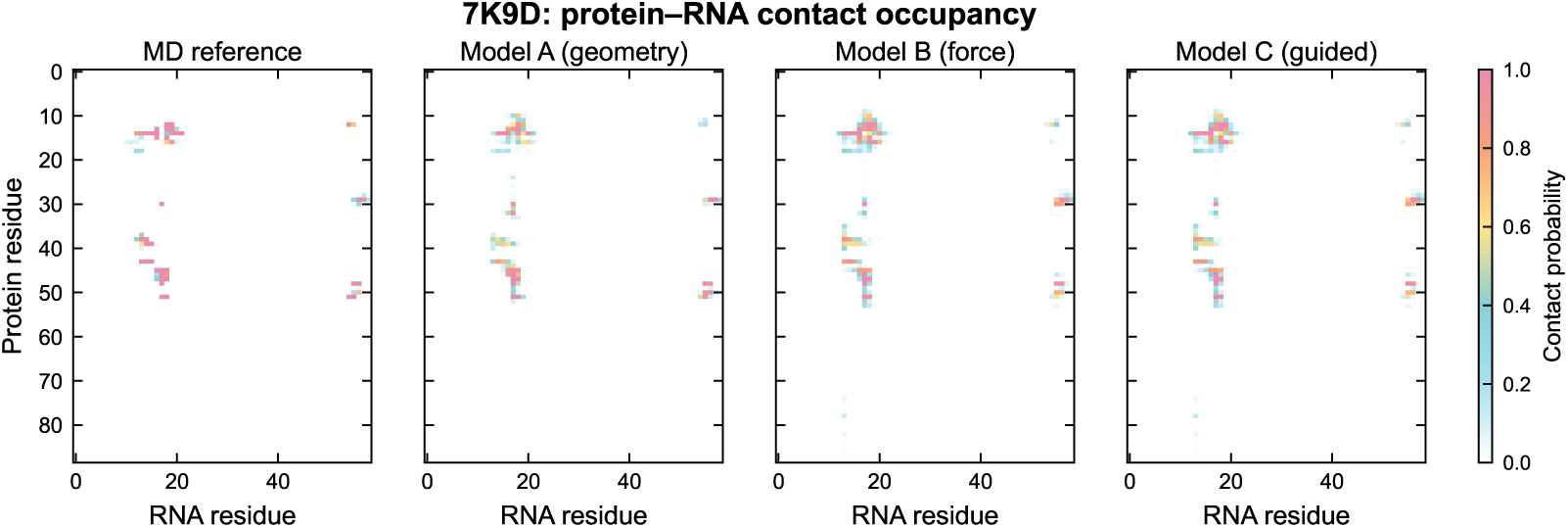
RNA–protein contact occupancy map for 7K9D. Layout follows the previous figure.

**Figure S7:**
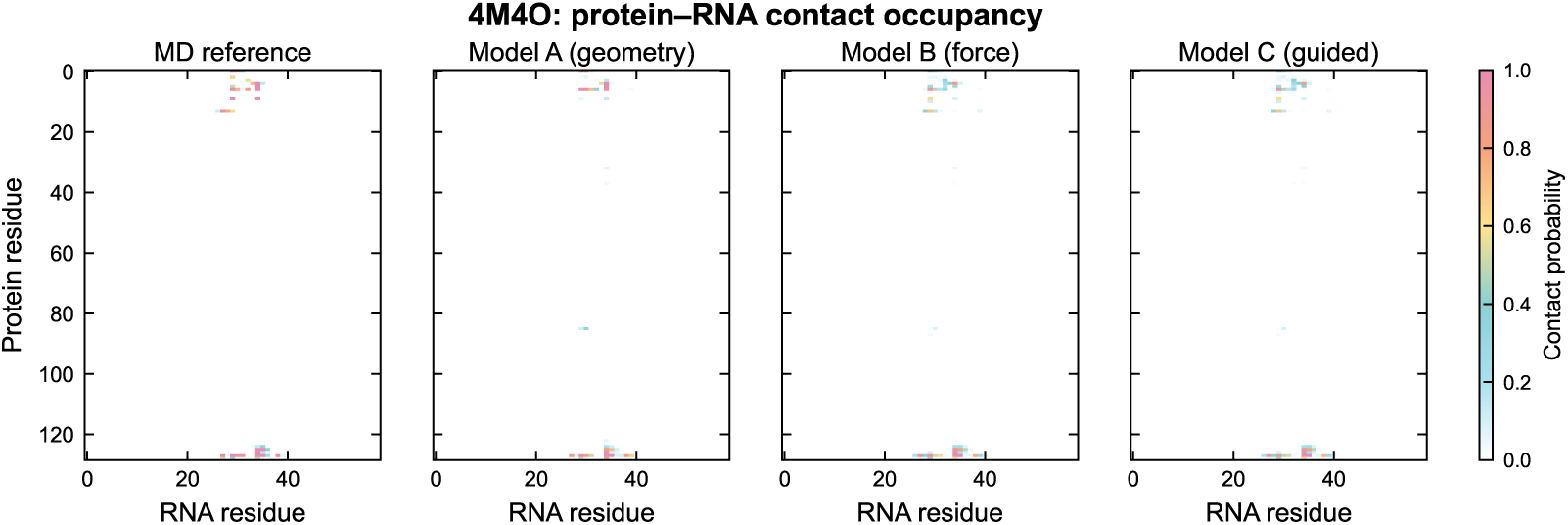
RNA–protein contact occupancy map for 4M4O. The largest test system shows more variable contact patterns in the MD reference, while generated ensembles maintain a more consistent contact profile.

**Figure S8:**
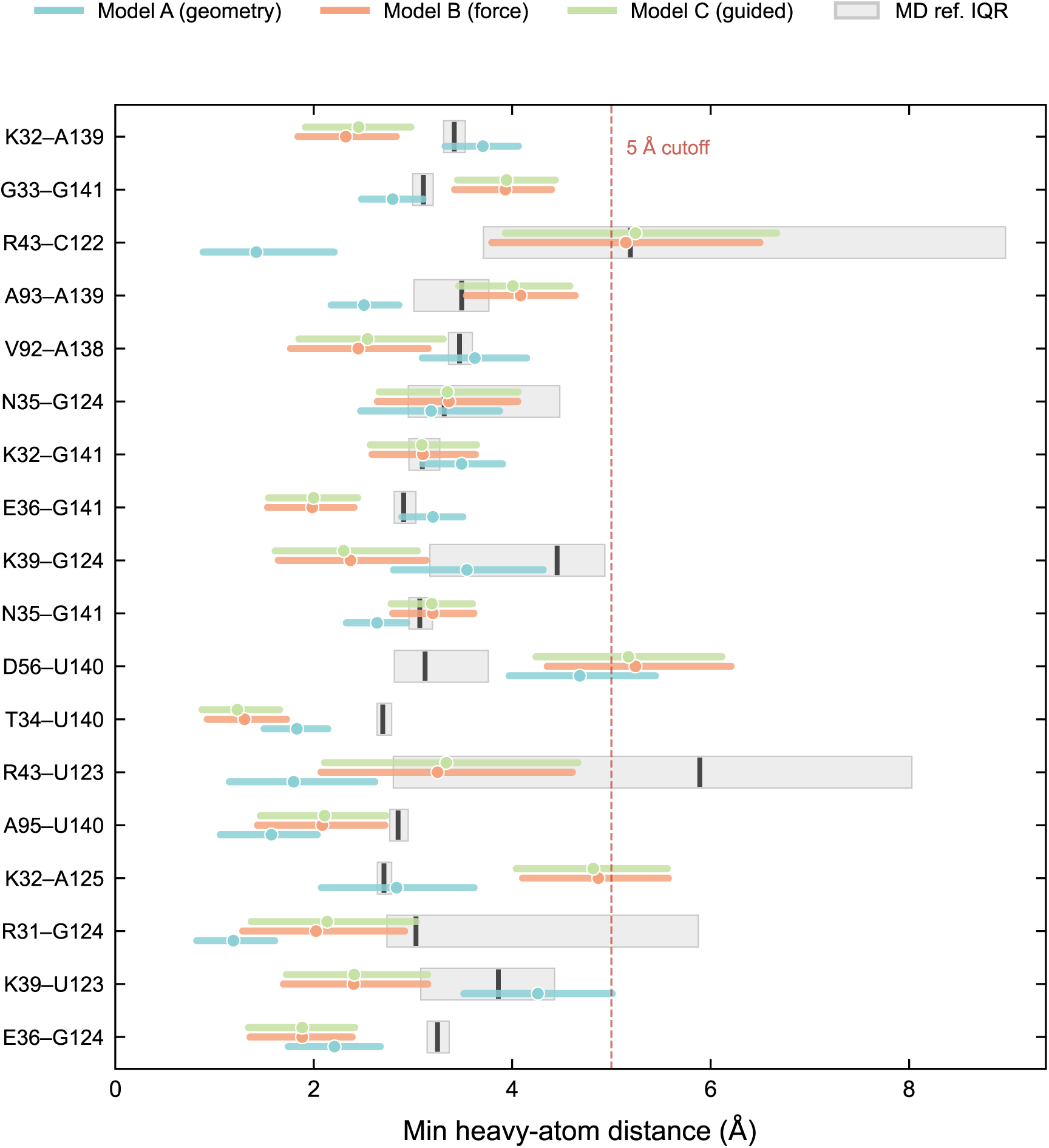
Per-contact-site distance distributions for 6TPH. Distributions follow the same layout as Figure 6.

**Figure S9:**
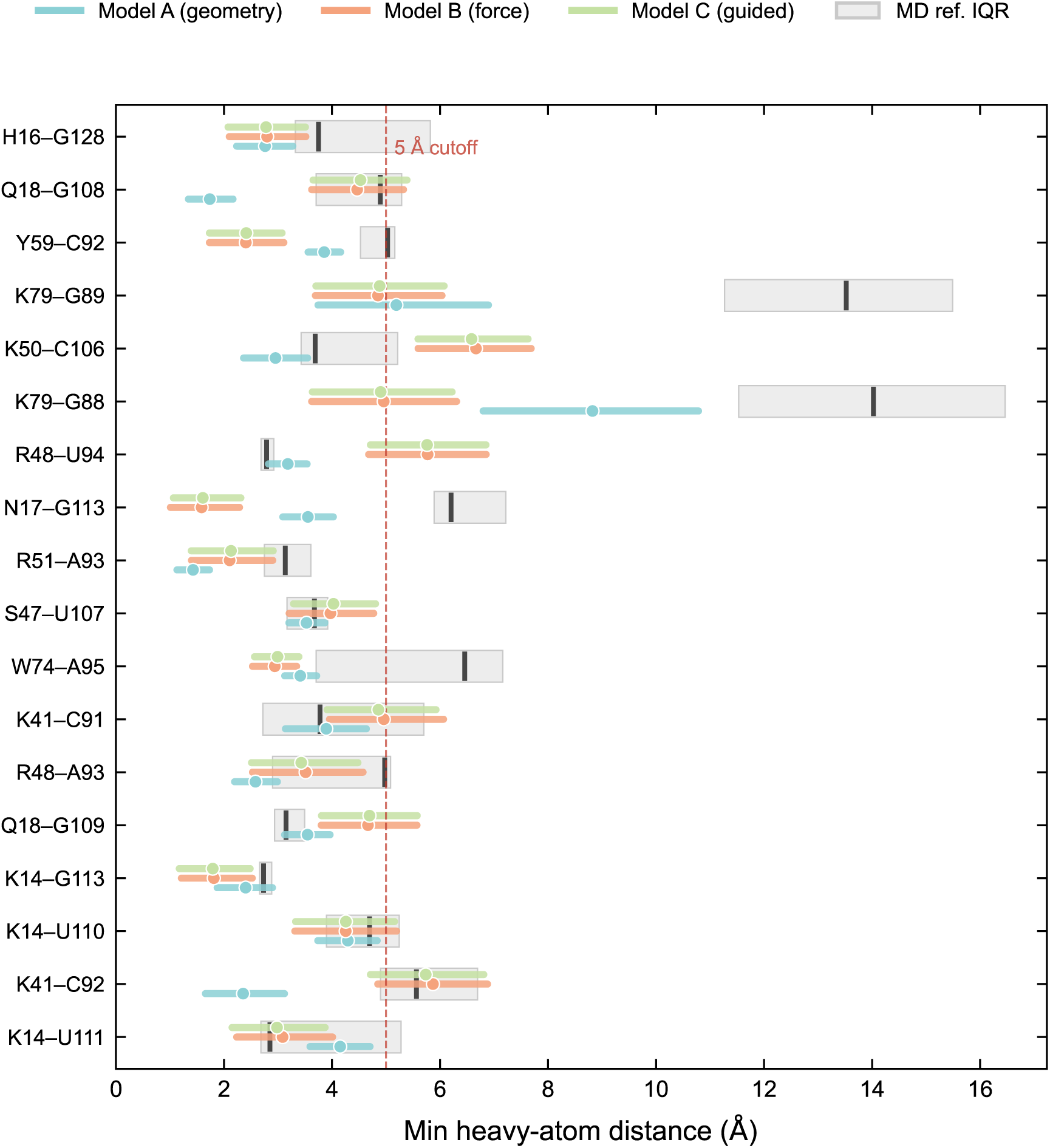
Per-contact-site distance distributions for 2HGH. The MD reference shows a broader distribution of interface distances than the generated ensembles.

**Figure S10:**
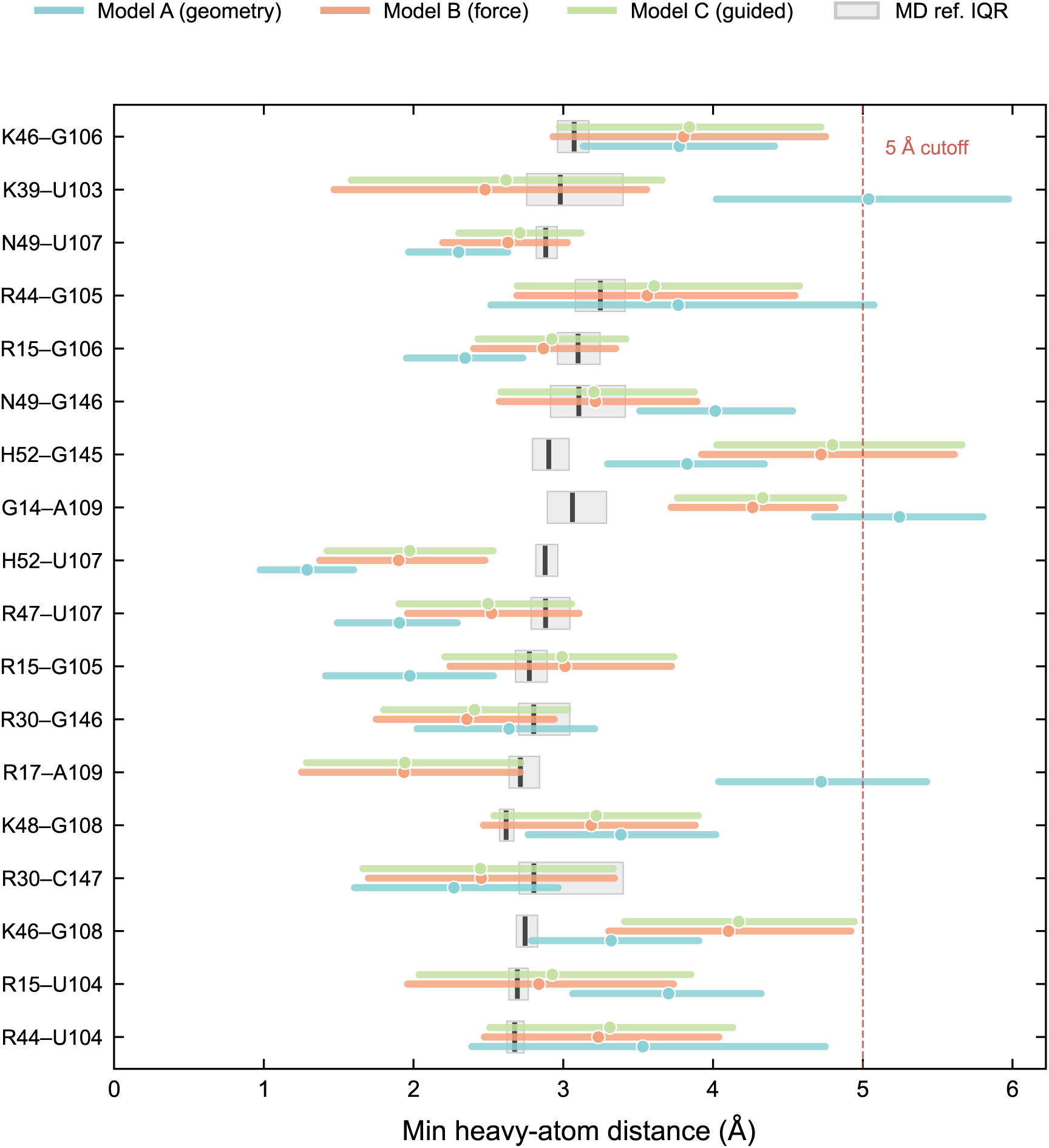
Per-contact-site distance distributions for 7K9D. Both MD and generated ensembles show moderate contact fluctuations.

**Figure S11:**
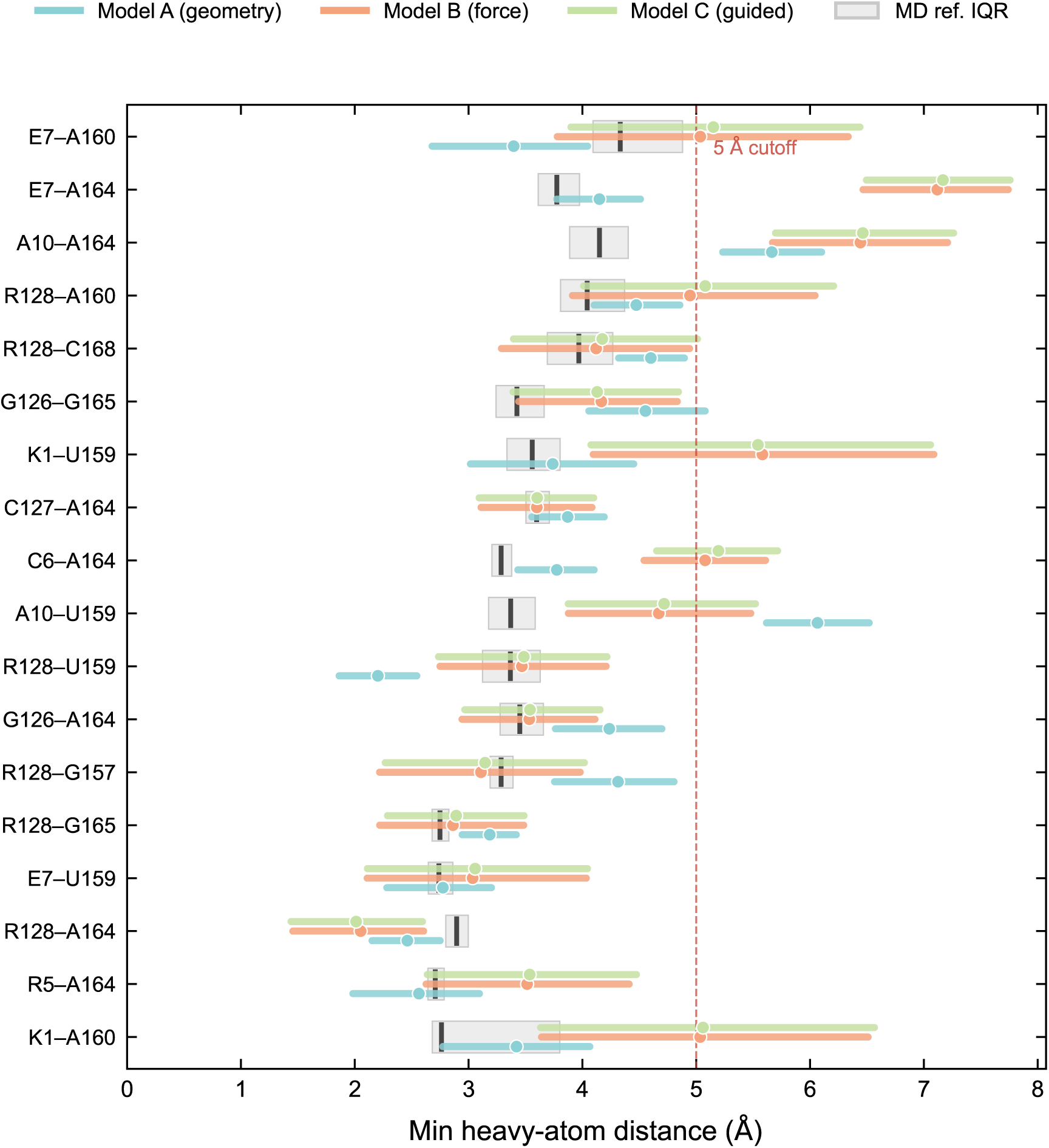
Per-contact-site distance distributions for 4M4O. The largest test system exhibits larger distance variations in the MD reference.

**Figure S12:**
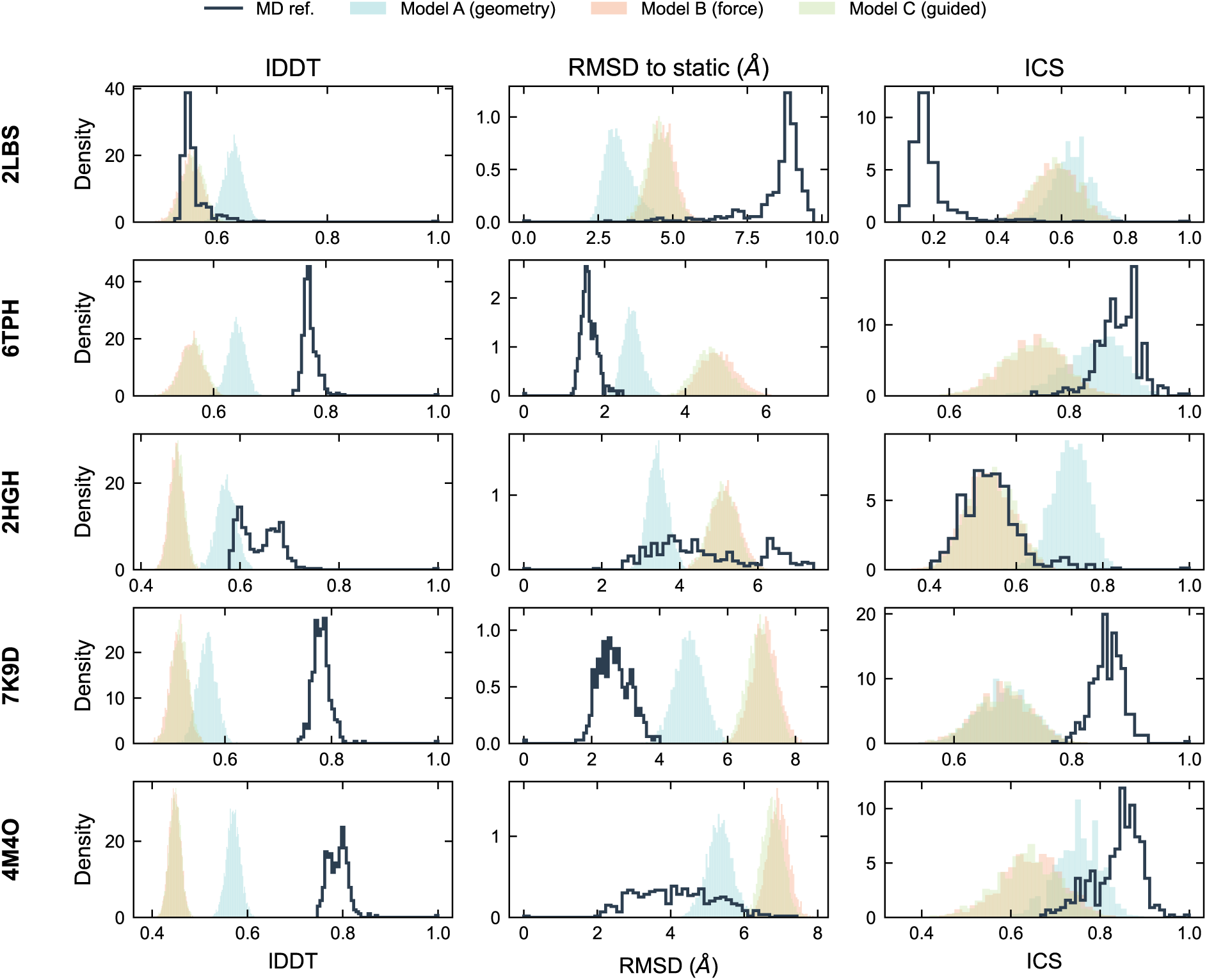
Per-conformation metric distributions across all test systems. Distributions of per-frame lDDT, RMSD to the static structure, and interface contact similarity are shown for all five test systems. Filled distributions correspond to generated ensembles, and outlined distributions to the MD reference. The generated distributions are more compact and centered near the static structure.

**Figure S13:**
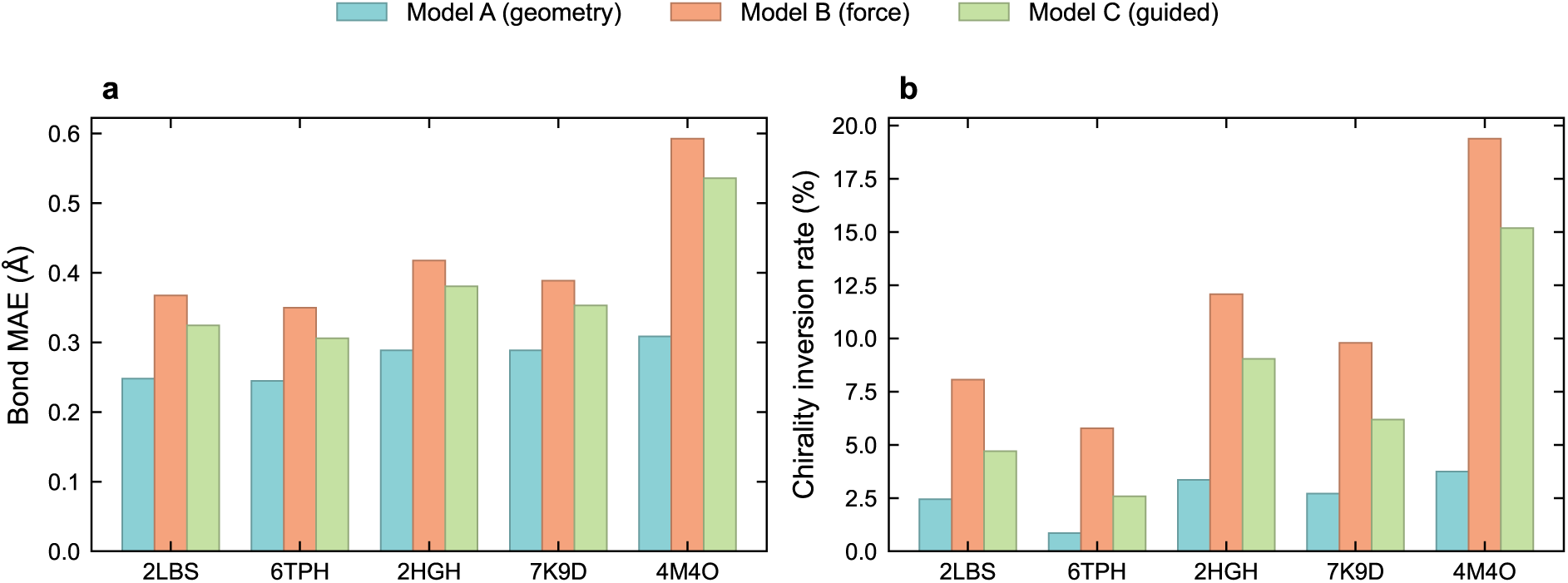
Chemical validity: bond-length mean absolute error and chirality inversion. Left: bond-length mean absolute error by system and model, partitioned into protein and RNA contributions. Right: chirality inversion rate. Model A maintains near-ideal geometry, while Models B and C show slightly elevated values, mainly in flexible loop regions.

**Figure S14:**
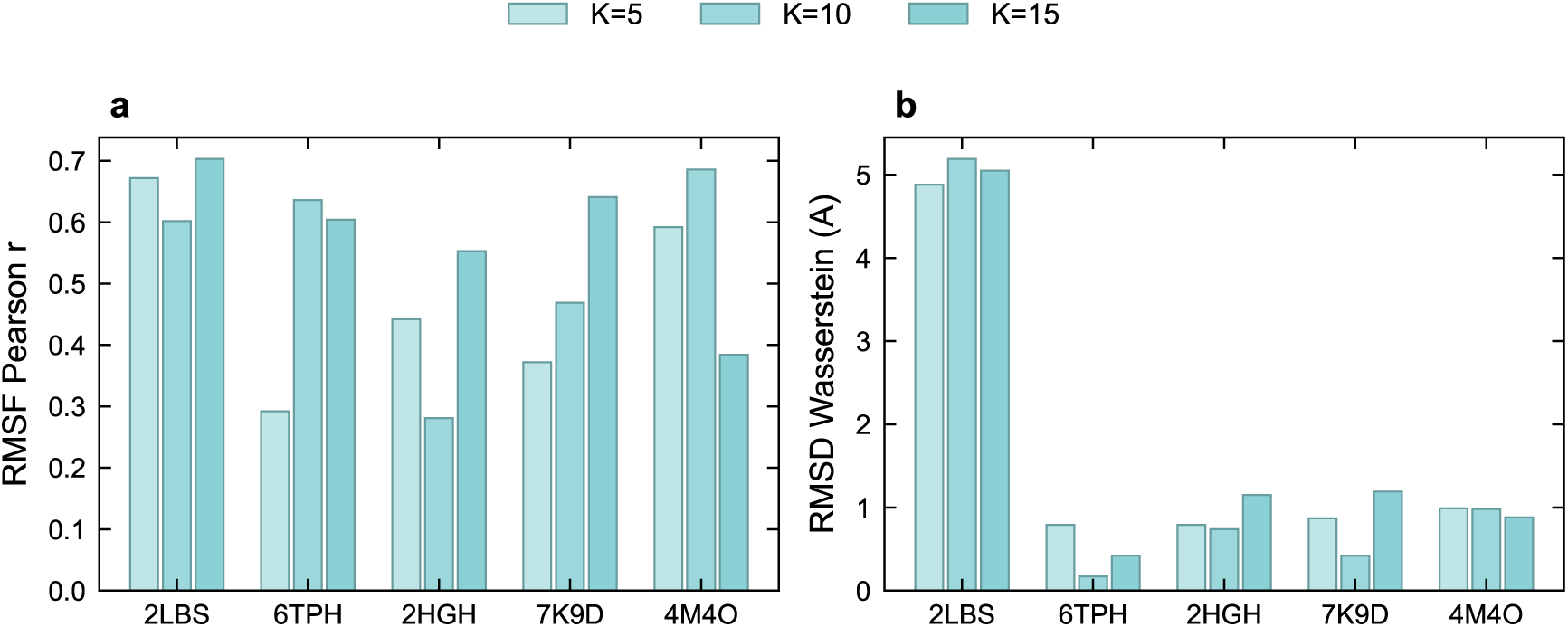
Data-scaling ablation. Model A checkpoints trained on different numbers of systems are evaluated on the five test systems. Performance improves with more training data, suggesting further gains are possible.

**Figure S15:**
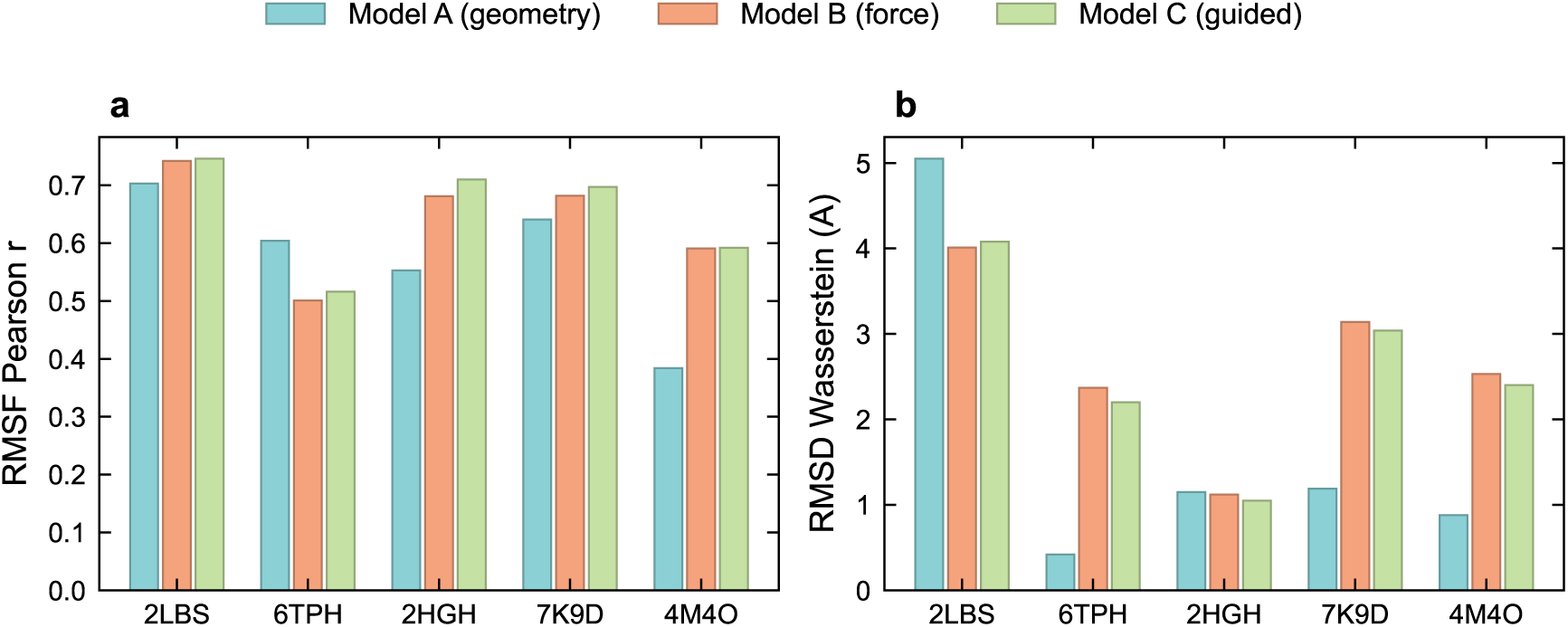
Independent benchmark metrics for all model variants. Per-system comparison of Models A, B, and C using the independent benchmark pipeline. The same relative trends observed in the primary evaluation are reproduced.

